# Synovial fibroblasts assume distinct functional identities and secrete R-spondin 2 to drive osteoarthritis

**DOI:** 10.1101/2022.05.06.489035

**Authors:** Alexander J. Knights, Easton C. Farrell, Olivia M. Ellis, Lindsey Lammlin, Lucas M. Junginger, Phillip M. Rzeczycki, Rachel F. Bergman, Rida Pervez, Monique Cruz, Alexa A. Samani, Chia-Lung Wu, Kurt D. Hankenson, Tristan Maerz

## Abstract

**Objectives:** Synovium is acutely affected following joint trauma and contributes to post-traumatic osteoarthritis (PTOA) progression. Little is known about discrete cell types and molecular mechanisms in PTOA synovium. We aimed to describe synovial cell populations and their dynamics in PTOA, with a focus on fibroblasts. We also sought to define mechanisms of synovial Wnt/β-catenin signaling, given its emerging importance in arthritis.

**Methods:** We subjected mice to non-invasive anterior cruciate ligament rupture as a model of human joint injury. We performed single-cell RNA-sequencing to assess synovial cell populations, subjected Wnt-GFP reporter mice to joint injury to study Wnt-active cells, and performed intra-articular injections of the Wnt agonist R-spondin 2 (Rspo2) to assess whether gain-of-function induced pathologies characteristic of PTOA. Lastly, we used cultured fibroblasts, macrophages, and chondrocytes to study how Rspo2 orchestrates crosstalk between joint cell types.

**Results:** We uncovered seven distinct functional subsets of synovial fibroblasts in healthy and injured synovium, and defined their temporal dynamics in early and established PTOA. Wnt/β-catenin signaling was overactive in PTOA synovium, and Rspo2 was strongly induced after injury and secreted exclusively by Prg4^hi^ lining fibroblasts. Trajectory analyses predicted that Prg4^hi^ lining fibroblasts arise from a pool of Dpp4+ mesenchymal progenitors in synovium, with SOX5 identified as a potential regulator of this emergence. We also showed that Rspo2 orchestrated pathological crosstalk between synovial fibroblasts, macrophages, and chondrocytes.

**Conclusions:** Synovial fibroblasts assume distinct functional identities during PTOA, and Prg4^hi^ lining fibroblasts secrete the Wnt agonist Rspo2 to drive pathological crosstalk in the joint after injury.

## INTRODUCTION

Inflammation, fibrosis, and mineralization of synovium are key pathological features of post-traumatic osteoarthritis (PTOA)(1-3). Despite major recent advances in our understanding of synovial biology in inflammatory conditions such as rheumatoid arthritis (4-7), little is known about mechanisms of PTOA pathogenesis.

Synovium is a heterogeneous connective tissue that exhibits intrinsic pathologies during PTOA and promotes disease throughout the joint(8). Synovial fibroblasts (SFs) are a mixed population of stromal cells that have been classified into pathological subsets in rheumatoid arthritis, based on their proliferative capacity, invasiveness, and localization, along with their ability to orchestrate inflammation or bone and cartilage damage(6, 7). However, in PTOA, functional SF subsets have not been defined beyond an anatomical delineation between Thy1+ sublining and Prg4^hi^ lining SFs(9). Given the strong focus on cartilage in osteoarthritis (OA), single-cell level analyses have shed significant light on chondrocyte subsets(10-12), but little attention has been given to disease-associated synovial cell populations.

Targeting overactive canonical Wnt/β-catenin signaling has emerged as a strong candidate for treating OA(13-16). However, the mediators and cell types responsible for overactive Wnt signaling in OA are not well described. R-spondin 2 (Rspo2) is a secreted, matricellular agonist of canonical Wnt signaling(17, 18), and recent reports have implicated its role in OA progression(19, 20).

Herein, to study synovial dynamics during PTOA, we employed a non-invasive anterior cruciate ligament rupture-based model of joint injury (ACLR) which recapitulates traumatic joint overloading and instability that occur in human joint injury without confounding effects from surgery(21-24). Transcriptomic profiling by single-cell RNA sequencing (scRNA-seq) uncovered distinct functional subsets of SFs in healthy and injured tissue. Further interrogation revealed overactive Wnt/β-catenin signaling throughout the SF niche in synovium, and pathological secretion of Rspo2 exclusively by Prg4^hi^ lining SFs. These insights offer new avenues for specific therapeutic targeting in PTOA.

## RESULTS

### Single-cell transcriptomic profiling of synovium reveals cellular heterogeneity in PTOA

The cellular composition of synovium has been characterized in inflammatory models of arthritis(6, 25), but little is known about synovial cell dynamics in PTOA. We used flow cytometry to demonstrate that synovial tissue digestion yielded viable (>90%) cells representing major resident subpopulations – synovial fibroblasts (SFs; CD31-CD45-), endothelial cells (CD31+ CD45-), and hematopoietic cells (CD31-CD45+) (Figure S1A). To study PTOA synovium, we employed a non-invasive, joint-overloading ACLR model, which induces key clinical features of PTOA including cartilage degradation, synovial inflammation, fibrosis, and osteophyte formation(22-24). We performed scRNA-seq on synovium combined from male and female mice 7d ACLR, 28d ACLR, or Sham controls (Figure S1B). After initial quality control, 20,422 cells were integrated across conditions by canonical correlation analysis(26) (Figure S1C-D), then unsupervised dimensionality reduction and clustering were performed (Figure 1A-B and Figure S2). SFs (sublining and lining), myeloid, and endothelial cells predominated numerically in healthy and PTOA synovium, with SFs and myeloid cells undergoing the most drastic increase in abundance at 7d ACLR (Figure 1C-D). We also detected rarer cell populations including pericytes, T cells, and Schwann cells. Profound injury-induced changes in synovial cellular composition were underpinned by changes in global transcriptomic profile among Sham, 7d, and 28d ACLR (Figure 1E). To examine intercellular communication between synovial cells, we used CellChat, which infers cell-cell signaling pathways based on curated interaction networks between ligands, receptors, agonists, and inhibitors(27). Outgoing and incoming communication patterns for each major synovial cell type were calculated based on relative contributions of ligand-receptor interactions, displayed as river plots (Figure 1F and Figure S3). These data demonstrate distinct, cell- and direction-specific communication signatures that orchestrate cell-cell signaling in synovium, allowing for subsequent analysis of networks of interest. Together these results highlight the plasticity of synovium following joint injury and shed light on the diverse cell types that populate PTOA synovium.

**Figure 1.**
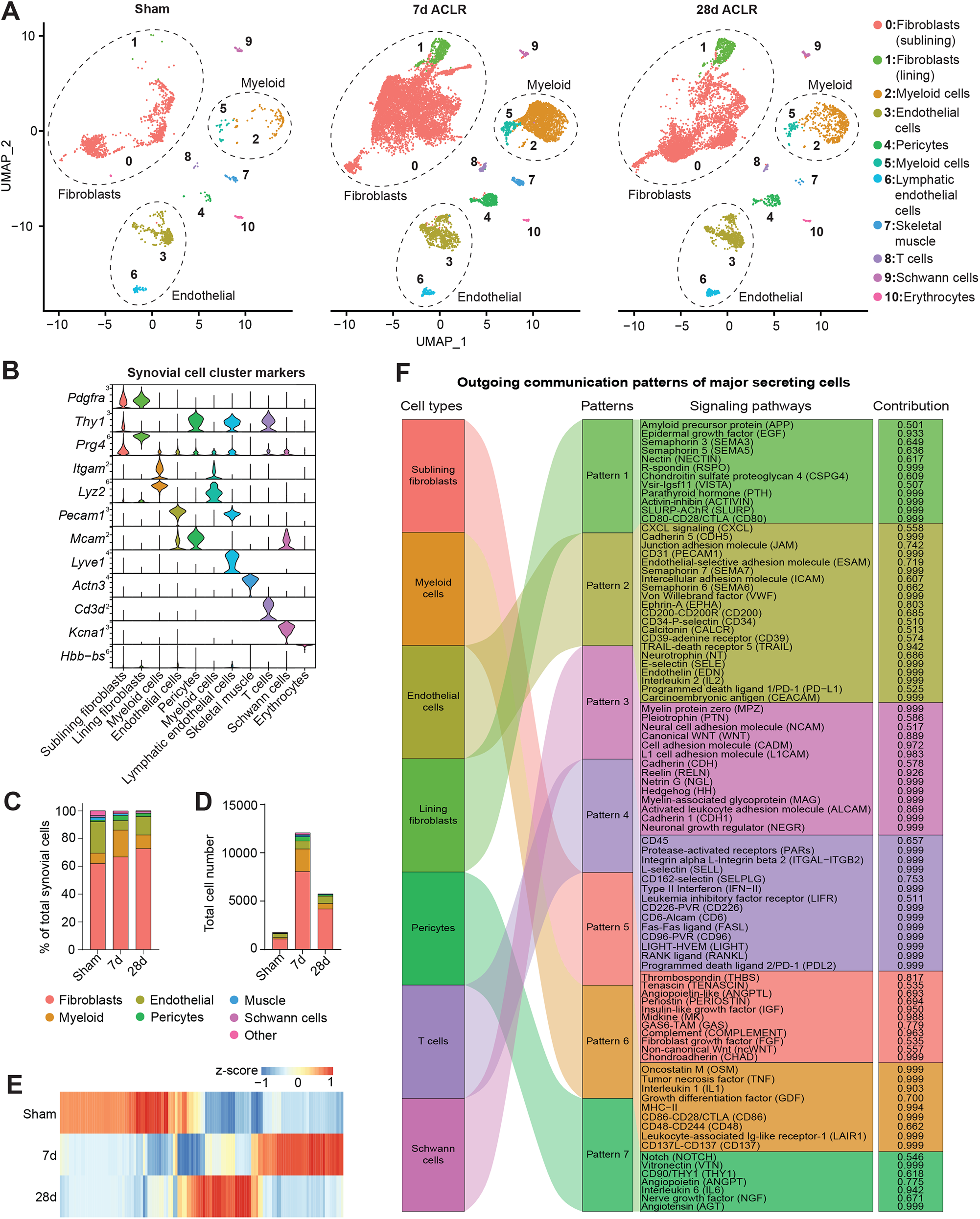
Single-cell transcriptomic profiling of synovium reveals cellular heterogeneity in PTOA. (A) UMAP plots showing reduced dimensionality projections of all synovial cells from Sham, 7d ACLR or 28d ACLR mice (20,422 cells, n=2 biological replicates per condition, each composed of one male and one female). Cell types are grouped by color and designated numbers, with annotations on the right. Major cell groups have dashed outlines. (B) Violin plots of synovial cell type marker genes. (C) Proportional breakdown of major synovial cell types in each condition. (D) Total abundance of major synovial cell types in each condition. (E) Global gene expression across conditions from pseudo-bulk analysis of scRNA-seq data of all synovial cells. (F) River plot from CellChat analysis showing outgoing signaling patterns from major synovial cell types and the pathways comprising each pattern (contribution score for each pathway is shown on the right). ACLR: anterior cruciate ligament rupture (injury model).

### Identification of functionally distinct synovial fibroblast subsets

For more granular interrogation of SF subpopulations, we computationally subset SFs and related mesenchymal cells, excluding pericytes. Re-clustering uncovered seven distinct subsets present in healthy synovium (Angptl7+, IL-6+, Dpp4+, Prg4^hi^) or induced after injury (αSMA+, Sox9+, Mki67+), each designated by highly confined expression of marker genes (Figure 2A-C and Figure S4). Established SF markers, such as *Vim, Pdgfra*, and *Pdpn* were broadly present across subsets, with a distinction between Thy1+ SFs (expressed in Angptl7+, αSMA+/Acta2+, IL-6+, Dpp4+, and Mki67+ clusters) and Prg4^hi^ Thy1-lining SFs (Figure 2D). Profound injury-induced changes in both proportion and abundance of subsets occurred by 7d ACLR; all clusters expanded numerically post-injury but the most robust increases were evident in subsets absent from healthy synovium – the αSMA+ and Sox9+ clusters (Figure 2E-G). Notably, the abundance of all subsets diminished by 28d ACLR, apart from Prg4^hi^ lining SFs, consistent with persistent lining hyperplasia observed in human PTOA(28, 29). Having defined distinct SF clusters and their temporal abundance, we assessed whether their unique transcriptomic profiles underpinned distinct functional roles. All clusters, except for Mki67+ cells, exhibited hallmark features of SFs, with roles in extracellular matrix and collagen organization (Figure S5A). Gene Ontology and Reactome analyses uncovered unique functions of each subset (Figure 2H and Figure S5B). Fibroblast subsets were also subjected to CellChat analysis to characterize broad signaling patterns (Figure 2I and Figure S5C-I). Angptl7+ cells, present in healthy and injured synovium, were enriched for pathways relating to neurogenesis and were important secretors of ligands for glial-derived neurotrophic factor signaling. Injury-induced αSMA+ cells were enriched for pathways including myofibril assembly, smooth muscle contraction, and response to mechanical stimulus – reminiscent of myofibroblasts. IL-6+ cells, present in Sham and expanded by injury, were enriched for pro-inflammatory and chemotactic signaling, suggesting a role in orchestrating immune-SF crosstalk. Dpp4+ cells, which expressed previously reported mesenchymal progenitor markers *Dpp4, Wnt2, Anxa3, Cd34* and *Pi16*(30, 31), were present both in Sham and ACLR. Prg4^hi^ SFs, which constitute the thin lining layer in healthy synovium that exhibits dramatic hyperplasia in OA, were enriched for cell junction regulation, R-spondin and syndecan signaling, and were overrepresented for incoming signals from other SF subsets. Sox9+ cells, which also express *Col2a1, Osx/Sp7*, and *Acan*, were enriched for chondrogenic and osteogenic pathways, along with glycolytic metabolism, consistent with an osteochondral progenitor (OCP) phenotype. These cells may give rise to osteophytes, bony growths that constitute an important clinical feature of PTOA(32). And lastly, Mki67+ cells exhibited a distinct proliferating phenotype, expressing known markers such as *Birc5* and *Top2a*. Together these data reveal the diversity of stromal subpopulations in healthy and PTOA synovium and point towards their distinct functions and signaling patterns.

**Figure 2.**
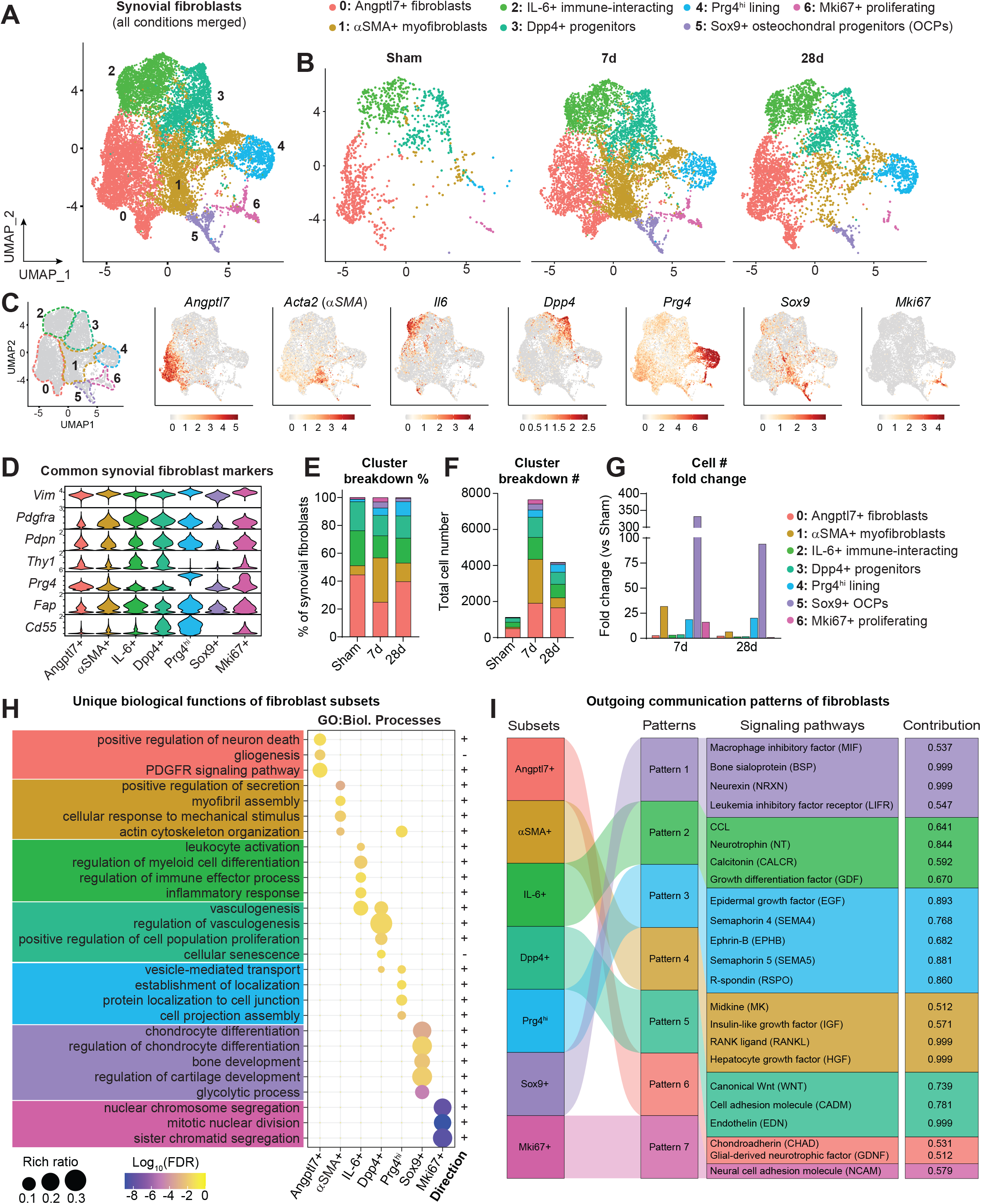
Identification of functionally distinct synovial fibroblast subsets. (A) UMAP plot showing SFs and related cell types integrated across conditions using canonical correlation analysis (n=2 biological replicates per condition, as in Figure 1). Clusters have designated numbers and colors corresponding to their annotation (top). (B) SF UMAP plots split by condition. (C) UMAP plot of SFs showing cluster borders with colored, dashed outlines (left), and feature plots of marker genes used to designate the naming of each cluster. Color scales represent relative expression for each gene individually. (D) Violin plots showing expression levels of common SF marker genes across clusters. (E) Proportional breakdown (%) of each cluster across condition. (F) Total abundance (#) of cells in each cluster across conditions. (G) Fold change (relative to Sham) of cell number (#) in each cluster at 7d and 28d ACLR. (H) Pathway analysis using Gene Ontology (GO) Biological Processes showing unique functional terms for each cluster and the direction of each term. (I) River plot showing CellChat analysis of outgoing signaling patterns from SF subsets and the pathways comprising each pattern, and the contribution score of each pathway on the right. FDR: false discovery rate.

### Canonical Wnt/β-catenin signaling is activated in synovium during PTOA

Targeting canonical Wnt/β-catenin signaling has therapeutic promise in OA given its reported overactivation in joint disease(14, 33, 34), and we found that the Wnt pathway was an important element of broad synovial signaling patterns in our study (Figure 1F and 2I). We showed Schwann cells to be important secretors of Wnt ligands in synovium, especially Wnt6 and Wnt9a, while Dpp4+ mesenchymal progenitors were important secretors within the SF niche (Figure 3A-B and Figure S6A-B). Beyond just outgoing Wnt signals, we showed that each of the SF subsets (except for Mki67+ proliferating SFs) were enriched for canonical Wnt signaling terms compared to non-SF cell types in synovium (Figure 3C). This observation is underpinned by upregulation of Wnt pathway genes in whole synovium and in only SFs, at 7d and 28d ACLR (Figure S6C-D). To assess synovial cells actively undergoing canonical Wnt/β-catenin signaling, we used Wnt-GFP reporter mice which express TCF/LEF response element-driven EGFP(35). The abundance, but not total proportion, of Wnt-GFP+ cells increased in 7d ACLR compared to contralateral synovium, and we observed increased GFP intensity – a proxy for increased Wnt/β-catenin signaling activity (Figure 3D-E and Figure S7A-B). Wnt-GFP+ cells were predominantly CD31-CD45-, indicative of SFs (Figure 3F), and total number and GFP intensity increased within this population in ACLR versus contralateral synovium (Figure 3G-H and Figure S7C). This corroborates our finding from scRNA-seq pathway analysis that SFs are the predominant Wnt-active cells in synovium (Figure 3C). These results demonstrate the importance of canonical Wnt/β-catenin signaling in synovial crosstalk and its overactivation in PTOA, with SFs being key cells undertaking Wnt signaling.

**Figure 3.**
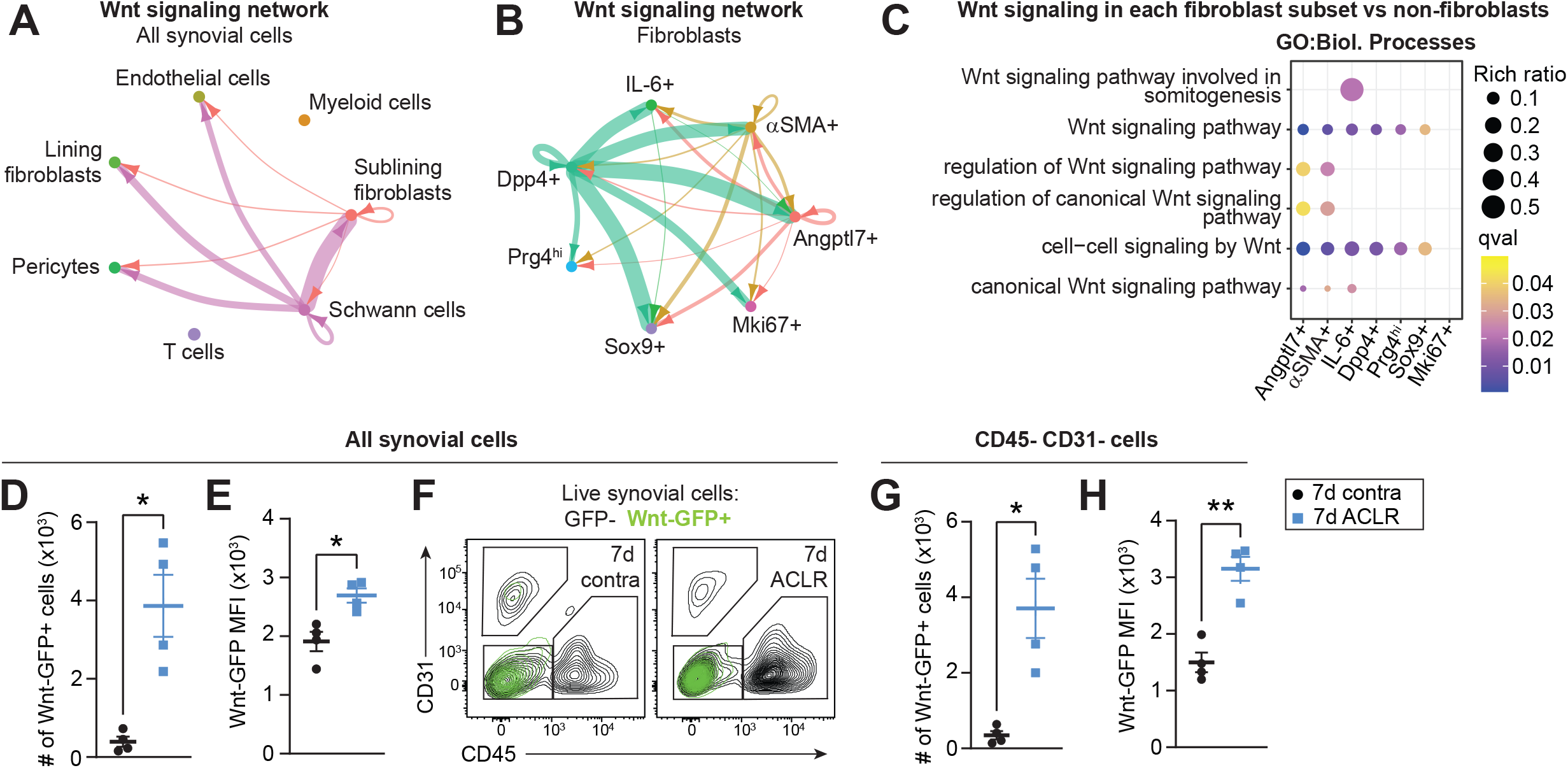
Canonical Wnt/β-catenin signaling is activated in synovium during PTOA. (A-B) Circle plots showing canonical Wnt signaling communication between all synovial cell types (A) or SFs only (B), with directionality indicated by arrowheads. Lines are color-coded by source cell type and line width is proportional to interaction strength. (C) Pathway analysis using GO Biological Processes showing enriched canonical Wnt signaling terms in each SF cluster compared to all non-SFs. (D-H) Flow cytometry assessment of injured or contralateral (contra) synovium from Wnt-GFP reporter mice 7d ACLR (n=4 mice). (D) Total number of Wnt-GFP+ synovial cells. (E) Median fluorescence intensity (MFI) of GFP for all synovial cells. (F) Overlay of Wnt-GFP+ cells (green) from contra and 7d ACLR synovium showing their expression of CD31 and CD45. CD31-CD45-cells represent predominantly SFs in synovium. (G) Total number of CD31-CD45-synovial cells that were Wnt-GFP+. (H) MFI of GFP for CD31-CD45-synovial cells. Paired two-tailed student’s t-tests were used for D, E, G, and H, where \**P*<0.05, \*\**P*<0.01. Error bars are mean ± SEM. qval: *P* value adjusted for false discovery rate.

### The canonical Wnt agonist R-spondin 2 is strongly and selectively induced during PTOA

To understand potential Wnt modulators activated during PTOA, we focused on R-spondin 2 (Rspo2), a secreted, matricellular agonist of canonical Wnt signaling that has been implicated in PTOA(19, 20). Immunostaining of whole joint sections from Sham and ACLR limbs revealed widespread up-regulation of Rspo2 in synovium and meniscus after injury, with strong expression evident in the osteophyte (Figure 4A-B). We detected elevated Rspo2 in synovial fluid, up-regulation of *Rspo2* transcript by qPCR, and increased abundance of RSPO2+ cells by flow cytometry, in synovium of injured mice (Figure 4C-D and Figure S8A). Despite a previous report that synovial M1 macrophages express Rspo2 in OA(20), we did not detect any CD45+ hematopoietic cells expressing Rspo2 by flow cytometry, in healthy or injured synovium, nor did scRNA-seq indicate *Rspo2* expression in myeloid cells (Figure 4E-F). *Rspo2* transcript and protein were only detected in SFs, where it was significantly more highly expressed than other R-spondin family members and strongly induced in PTOA (Figure 4E-F and Figure S8B-D). This observation was consistent in primary cultured cells, where *Rspo2* was highly expressed in SFs compared to bone marrow-derived macrophages (BMDM) (Figure S8E-F). Within SFs, Rspo2 was confined to the Prg4^hi^ lining subset, not Thy1+ sublining SFs (Figure 4G), consistent with Rspo2 immunostaining localization to the synovial lining (Figure 4A). Cell-cell signaling analysis revealed that outgoing R-spondin signaling was dominated by interactions between lining-derived Rspo2 and its receptors Lgr4, 5, and 6 on other SF subsets (Figure 4H and Figure S8G). To assess the injury-associated emergence of Prg4^hi^ Rspo2-expressing lining SFs, we performed trajectory analysis within the SF niche using Monocle3. This analysis identified Dpp4+ progenitors as precursors for Prg4^hi^ lining SFs across pseudotime, via an αSMA+ intermediate akin to that seen in other musculoskeletal injury models such as fracture healing(36, 37) (Figure 4I and Figure S8H). This differentiation was supported by pseudotime regression analysis of genes unique to the Dpp4+ root (*Dpp4, Pi16, Wnt2*) or the Prg4^hi^ terminus (*Prg4, Col22a1, Rspo2*) of the proposed trajectory (Figure 4J-K). Transcription factor motif analysis of genes derived from co-regulation network analysis along the trajectory revealed SOX5 as a putative regulator of *Rspo2* in lining SFs (Figure 4L-N and Figure S8I). Together these results demonstrate that Rspo2 is induced in PTOA and, within synovium, is exclusively secreted by lining SFs. Prg4^hi^ lining SFs that secrete Rspo2 arise from a Dpp4+ progenitor pool within synovium, and this differentiation trajectory, regulated in part by SOX5, may provide insights into mechanisms of chronic lining hyperplasia in PTOA.

**Figure 4.**
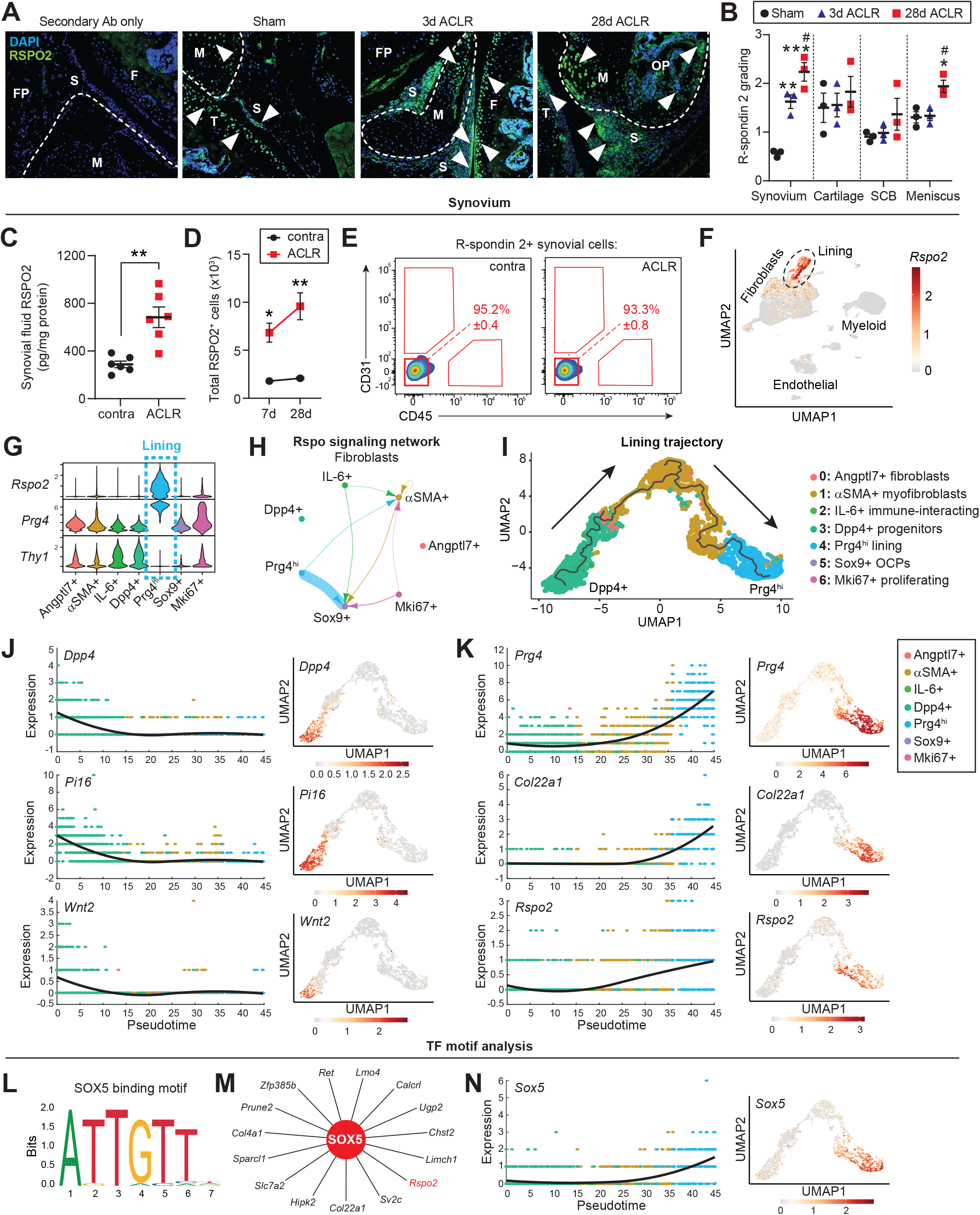
The canonical Wnt agonist R-spondin 2 is strongly and selectively induced during PTOA. (A) Immunofluorescent staining of R-spondin 2 (RSPO2) in joint sections from Sham, 3d or 28d ACLR. A negative control with only secondary antibody (Ab) staining is included (left). Nuclei were counterstained with DAPI. White arrowheads indicate areas of RSPO2 expression. Representative images are shown from n=3 mice per condition. FP: fat pad; S: synovium; F: femur; T: tibia; M: meniscus; OP: osteophyte. (B) Qualitative grading of RSPO2 staining in (A) for synovium, cartilage, subchondral bone (SCB), and meniscus. (C) RSPO2 protein levels in synovial fluid from contralateral (contra) and 28d ACLR (n=6 mice), measured by ELISA. Amount of RSPO2 was normalized to total protein in each synovial fluid sample. (D-E) Assessment of RSPO2+ cells from contra, 7d ACLR or 28d ACLR synovium by flow cytometry (n=3 mice per condition). (D) Total number of RSPO2+ cells in synovium. (E) Expression of CD31 and CD45 in RSPO2+ synovial cells. Mean percentage of RSPO2+ cells in the CD31-CD45-gate is shown in red ± SEM. (F) Feature plot showing expression of *Rspo2* in all synovial cells by scRNA-seq. Lining SFs are outlined. (G) Violin plots of *Rspo2*, the sublining SF marker *Thy1*, and the lining marker *Prg4* across SF subsets. (H) Circle plot showing Rspo signaling communication between SF subsets, with directionality indicated by arrowheads. Lines are color-coded by source cell type and line width is proportional to interaction strength. (I) Trajectory analysis of Prg4^hi^ lining SFs. The black line represents the trajectory calculated by Monocle3 and cells are color-coded according to their cluster. Arrows indicate progression of pseudotime derived from Figure S7H. (J-K) Pseudotime regression plots (left) and the corresponding feature plots mapped onto the lining trajectory UMAP (right), for genes associated with the (J) Dpp4+ root (*Dpp4, Pi16, Wnt2*) or (K) Prg4^hi^ terminus (*Prg4, Col22a1, Rspo2*) of the trajectory. (L-N) Transcription factor motif analysis in Module 2 (from Figure S8H) of the Prg4^hi^ lining trajectory, using RcisTarget. (L) Binding motif for SOX5 (jaspar__MA0087.1). (M) Gene regulatory network for SOX5, generated by CytoScape. (N) Pseudotime regression plot (left) and feature plot (right) for *Sox5* in the Dpp4+ to Prg4^hi^ trajectory. For (B), one-way ANOVA with multiple comparisons and Tukey’s post-hoc correction was used to assess significance in each tissue compartment (\**P*<0.05, \*\**P*<0.01, \*\*\**P*<0.001 compared to Sham; #*P*<0.05 compared to 3d ACLR). For (C), a paired two-tailed student’s t-test was used with \*\**P*<0.01. For (D), a two-way ANOVA with multiple comparisons and Tukey’s post-hoc correction was used, where \**P*<0.05, \*\**P*<0.01 compared to the corresponding contralateral. For B-D, error bars are mean ± SEM.

### R-spondin 2 is sufficient to induce pathology in healthy joints of mice

To study gain-of-function effects *in vivo*, we performed intra-articular injections of RSPO2 (right hindlimbs) or PBS (left hindlimbs) into mice and assessed knee hyperalgesia, matrix metalloproteinase (MMP) activity, and histological features of OA and synovitis (Figure 5A). Limbs injected with RSPO2 had a lower pain threshold, indicative of RSPO2-induced hyperalgesia, and higher MMP activity compared to PBS-injected limbs (Figure 5B-C). RSPO2-injected joints had elevated PTOA and synovitis severity scores, driven by proteoglycan loss in articular cartilage, synovial lining hyperplasia, fibrosis, inflammatory cell infiltrate, and cortical bone erosion (Figure 5D-G), clearly demonstrating that exogenous delivery of RSPO2 is sufficient to induce pathological features of PTOA in healthy mouse joints.

**Figure 5.**
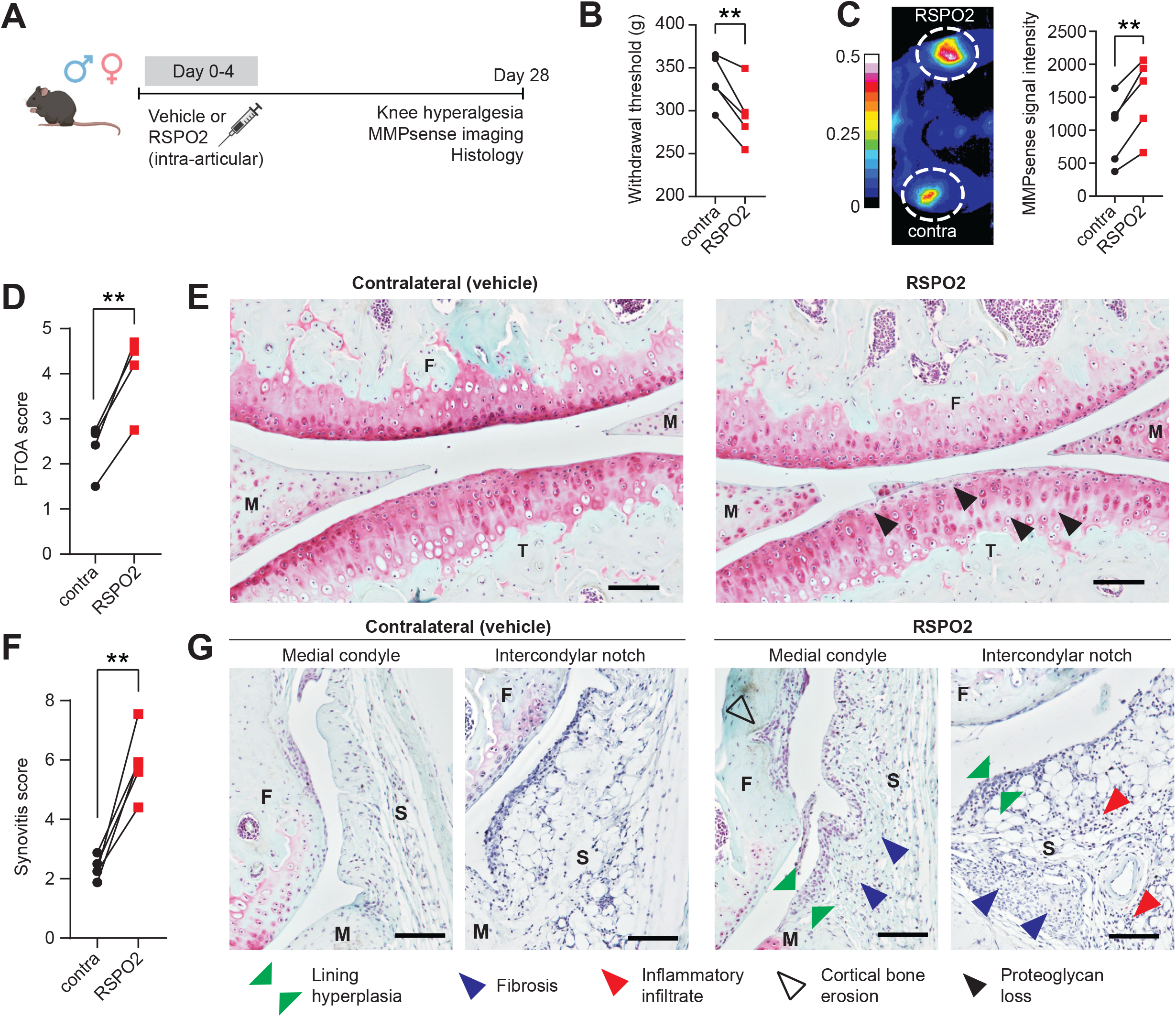
R-spondin 2 is sufficient to induce pathology in healthy joints of mice. (A) Experimental design. Male and female C57Bl6/J mice (n=5) were given intra-articular injections with R-spondin 2 (RSPO2, 200 ng/mL, right limb) or PBS (left limb) for five consecutive days. 28 days later, knee hyperalgesia and matrix metalloproteinase (MMP) activity were assessed, then whole joint histological sections were graded for OA severity and synovitis. (B) Knee hyperalgesia testing of contralateral (contra, PBS-treated) or RSPO2-treated limbs 28d after first injection. Paired limbs are connected by lines. (C) MMP activity as measured by near-infrared imaging of MMPSense680 probe. Representative intensity image (left) and quantitation of signal (right). Paired limbs are connected by lines. (D-G) PTOA (D) and synovitis (F) severity scoring(49, 50) for contralateral versus RSPO2-injected limbs, with paired limbs connected by lines. Safranin O/Fast Green stained sections imaged at 10x magnification that are representative of (E) PTOA score or (G) synovitis score. Highlighted features are proteoglycan and matrix loss (black arrows), synovial lining hyperplasia (green arrows), synovial fibrosis (blue arrows), sub-synovial inflammatory infiltrate (red arrows), and cortical bone erosion (black outline arrows). Scale bar: 100 μm. For B, C, D and F, paired two-tailed student’s t-tests were used, where \*\**P*<0.01. F: femur; T: tibia; M: meniscus; S: synovium.

### R-spondin 2 orchestrates pathological crosstalk between joint-resident cell types

The joint is a complex organ with diverse cell and tissue types capable of signaling to one another via synovial fluid. Having shown that SFs produce Rspo2 and that it is enriched in synovial fluid of injured joints, we sought to assess how SF-derived Rspo2 interacts with other joint-resident cell types. Treatment of SFs with inflammatory cytokines TNFα or IL-1β, known to be activated in OA(38, 39), did not increase secretion of Rspo2 into conditioned media (Figure 6A). *In vivo*, SFs expressed *Lgr4, Lgr5* and *Lgr6*, the receptors for Rspo2; *Lgr4* was broadly expressed by SFs, whereas *Lgr5* and *Lgr6* were confined to the αSMA+ and IL-6+, and the Sox9+ clusters, respectively, and are induced after injury (Figure 6B and Figure S9A). SFs derived from murine hindpaws or knees expressed high levels of *Lgr4* and *Lgr5*, thus serving as valid models for studying Rspo2 signaling in SFs (Figure S9B). Treatment of Wnt-GFP reporter-derived SFs with RSPO2 increased the percentage of Wnt-GFP+ cells and GFP intensity (Figure 6C). We observed induction of Wnt-responsive genes *Axin2* and *Lef1*, and of *Rspo2* itself, after treating SFs with RSPO2 (Figure 6D). This induction was mitigated by treatment with the Lgr inhibitor Mianserin(19). These data demonstrate that SFs secrete and respond to RSPO2, at least in part via Lgr signaling. Macrophages, on the other hand, did not express Lgr receptors (Figure S9D). We saw no induction of Wnt genes (*Axin2, Lef1*) or macrophage functional genes (*Il1b, Tnf, Mrc1/Cd206, Il10*) upon treatment of BMDM with RSPO2 (Figure 6E). Therefore, we treated BMDM with conditioned media from vehicle-or RSPO2-treated SFs, to ask whether SFs secrete secondary mediators that affect macrophage polarization. SF conditioned media was given to BMDM polarized towards M1 (IL-1β), M2 (IL-4) or M0 (PBS) (Figure 6F). We observed that in each condition, media from RSPO2-treated SFs skewed macrophage polarization towards an inflammatory, M1-like phenotype, based on composite gene expression of M1/M2 markers (Figure 6G). This was mitigated when SFs were co-treated with Mianserin, reinforcing that RSPO2-induced activation of SF-derived secondary mediators that activate macrophages is Lgr-dependent. We then employed this conditioned media treatment protocol to chondrogenic ATDC5 cells, which express Lgr receptors (Figure S10A). Treatment of differentiated ATDC5 cells with media from RSPO2-treated SFs increased their expression of osteogenic genes *Bsp* and *Ocn* (Figure S10B). We also showed that direct treatment of ATDC5 cells with RSPO2 during and after chondrogenic differentiation suppressed chondrogenic genes, but elevated hypertrophic, osteogenic, and Wnt genes (Figure S10C-D), consistent with previous reports of a pro-osteogenic and anti-chondrogenic role for Rspo2(19, 40). Together, these results demonstrate that the mechanism by which Rspo2 orchestrates pathological crosstalk in the joint involves pro-inflammatory signaling between SFs and macrophages, and pro-hypertrophic signaling between SFs and chondrocytes.

**Figure 6.**
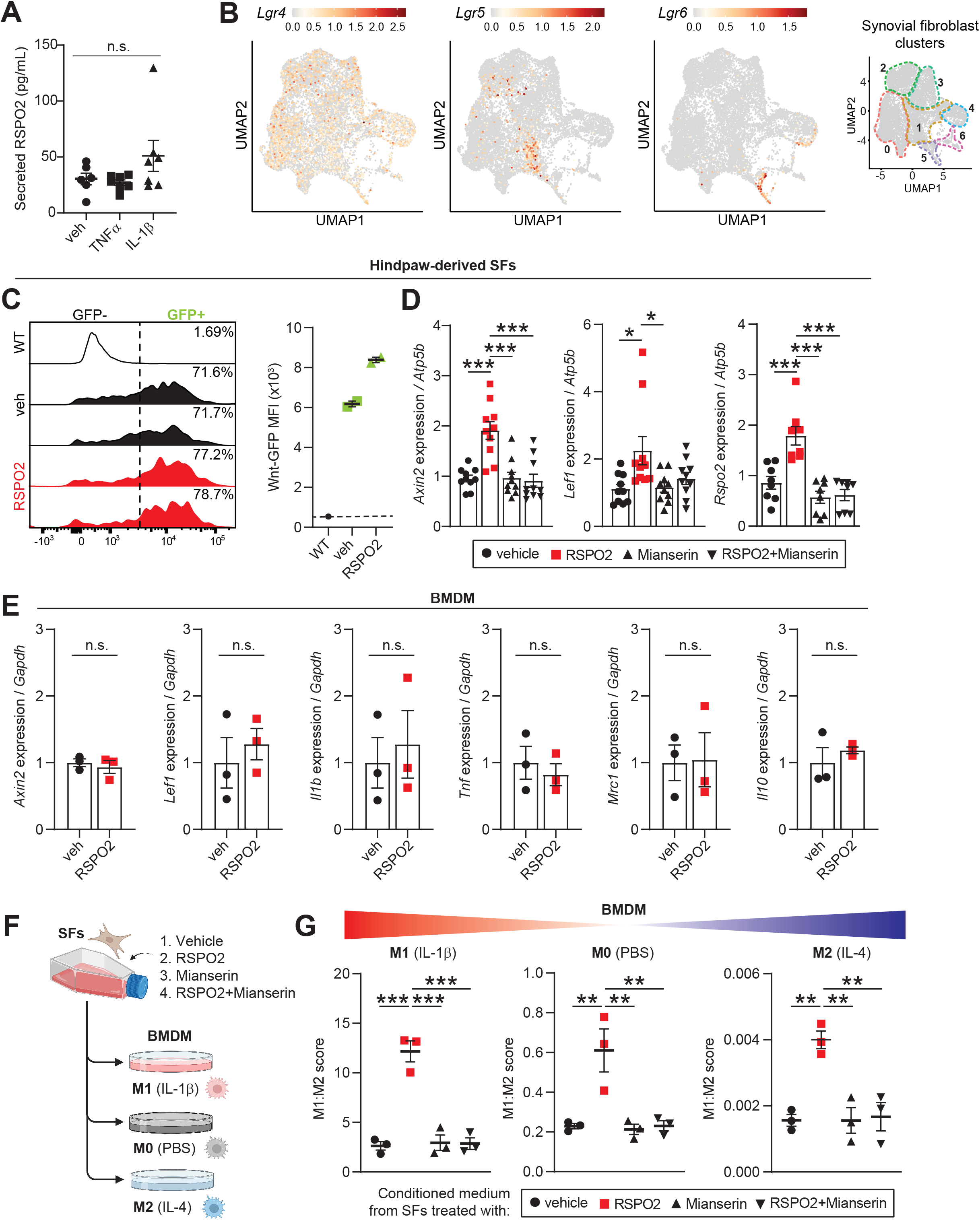
R-spondin 2 orchestrates pathological crosstalk between joint-resident cell types. (A) Secreted RSPO2 protein levels in conditioned medium from cultured SFs treated for 48 h with vehicle (veh), TNFα (10 ng/mL) or IL-1β (10 ng/mL). (B) Feature plots of Lgr receptors (*Lgr4-6*) in SFs. Color scales are not equivalent between plots. Cluster borders are shown on the right. (C) Hindpaw-derived SFs from Wnt-GFP reporter mice were treated with veh or RSPO2 (200 ng/mL) for 24 h. SFs were analyzed by flow cytometry to assess the total percentage of Wnt-GFP+ cells (left) and Wnt signaling activity (Wnt-GFP MFI, right). WT SFs (no reporter) were used as a negative control. Two biological replicates were performed. (D) Hindpaw-derived SFs were treated for 24 h with veh, RPSO2 (200 ng/mL), Mianserin (20 μM), or RSPO2+Mianserin (n=8-10 biological replicates). Expression of *Axin2, Lef1* and *Rspo2* was normalized to *Atp5b* levels, and veh samples were set to 1. (E) Bone marrow-derived macrophages (BMDM) were treated for 8 h with veh or RSPO2 (200 ng/mL) (n=3 biological replicates). Expression of *Axin2, Lef1, Il1b, Tnf, Mrc1* and *Il10* were normalized to *Gapdh* levels, and veh samples were set to 1. (F) Schematic showing experimental design of SF-BMDM conditioned media treatment in (G). (G) Hindpaw-derived SFs were treated with vehicle, RPSO2 (200 ng/mL), Mianserin (20 μM), or RSPO2+Mianserin. After 6 h, treatment media was removed, cells were washed and replenished with normal SF media for another 18 h (24 h total). SF conditioned media was harvested and directly treated to BMDM for 8 h in tandem with M0 polarization (PBS, middle), M1 polarization (10 ng/mL IL-1β, left), or M2 polarization (10 ng/mL IL-4, right) (n=3 biological replicates per condition). A composite M1:M2 polarization score was calculated based on the expression of M1 (*Il1b, Il6, Nos2*) and M2 (*Mrc1, Il10, Arg1*) genes by qPCR. *Gapdh* was used as the housekeeper gene. For A, D and G, one-way ANOVA with multiple comparisons and Tukey’s post hoc testing was performed where \**P*<0.05, \*\**P*<0.01, \*\*\**P*<0.001. For E, paired two-tailed student’s t-tests were performed. Error bars are mean ± SEM. n.s.: not significant.

## DISCUSSION

Synovium undergoes dramatic changes in composition and function after joint injury; however, underlying molecular mechanisms are not well understood. Here, we report a hierarchy of functionally distinct SF subsets in healthy and injured joints, and demonstrate that lining SFs secrete the Wnt/β-catenin agonist R-spondin 2, which promotes PTOA.

Recent studies utilizing ‘omics approaches have shed light on SF and macrophage subpopulations in rheumatoid arthritis synovium, in mice and humans(5-7, 25, 41). Single-cell studies of OA, in addition to their paucity, have largely focused on cartilage(9-12, 42), given its historical and prevailing position as the focal point of OA research. Reports are now emerging focused on the role of synovium specifically in OA, including recent evidence from Nanus *et al* showing a SF gene signature of inflammation and neurotrophic factors linked to early OA pain(43). Studies in human(9) and mouse(10) defined the broad distinction between sublining and lining SFs in OA. Here, we have extended these findings to propose distinct functional subsets within the SF niche, including Dpp4+ mesenchymal progenitors and IL-6+ immune-interacting SFs, along with αSMA+ myofibroblast-like cells and osteochondral progenitors that arise after injury. These functional subsets are worthy of future *in vitro* and *in vivo* characterization.

Based on human GWAS evidence linking canonical Wnt/β-catenin signaling to OA(44-46), ongoing pre-clinical trials aim to harness broad inhibition of Wnt signaling for OA treatment(15, 16). R-spondins are potent Wnt agonists that facilitate Frizzled/LRP receptor stability to permit Wnt activation(18). Okura *et al* showed that Rspo2 is elevated in synovial fluid of OA patients(19), and identified the antidepressant Mianserin as a putative Rspo/Lgr inhibitor that mitigated anti-chondrogenic actions of Rspo2. Another study showed that intra-articular injection of RSPO2 exacerbated collagenase-induced OA in mice, consistent with our results in uninjured mice, and demonstrated efficacy of anti-Rspo2 treatment in suppressing β-catenin overactivation(20). The authors, however, proposed that Rspo2 was secreted by synovial macrophages, for which we found no evidence in healthy or injured synovium, nor *in vitro*. Instead, we used multiple approaches to demonstrate that Prg4^hi^ lining SFs express and secrete Rspo2. Differentiation trajectory analysis pointed towards a Dpp4+ mesenchymal progenitor pool giving rise to Rspo2-expressing lining SFs, via an αSMA+ intermediate. Advances in multiomic single-nucleus sequencing, combining gene expression and chromatin accessibility for refined trajectory inference(47), will facilitate further interrogation of lining SF differentiation and provide insights into pathological lining hyperplasia seen in PTOA. Given widespread induction of Rspo2 that we observed throughout the joint, further work examining how Rspo2 drives PTOA-relevant pathologies such as fibrosis, osteophytes, and inflammation is warranted. Building on promising findings of Rspo/Lgr inhibition by Mianserin, the development of specific and sustained-release OA therapies that target overactive Wnt and Rspo signaling should also be prioritized.

The lack of longitudinal samples and bias towards end-stage donors are inherent drawbacks of human tissue samples, complicating studies focused on early disease etiology. Therefore, the clinically-relevant ACLR model of PTOA in mice provides a useful means of studying pathophysiological events that precede established disease. Nevertheless, it will be critical to assess the extent to which our findings, especially distinct SF subsets, are conserved in human PTOA. Changes in gene expression and preferential cell type depletion are shortcomings of common tissue dissociation methods, as used here. The rapid advancement of spatial transcriptomics may remedy this concern and allow scientists to capture both the phenotypic and anatomical dimension, which is essential for an organ as architecturally complex as the joint. Meta-analysis of both mouse and human single-cell studies from OA synovium and cartilage will further help to bridge gaps in knowledge, harmonizing existing data for better utility and interpretation.

In summary, we defined distinct functional subsets of SFs in PTOA, showed that lining SFs secrete Rspo2, and that Rspo2 promotes pathological crosstalk within the joint. We also identified a Dpp4+ progenitor pool that gives rise to Prg4^hi^ Rspo2-expressing lining SFs. These findings highlight the importance of SF speciation and canonical Wnt/β-catenin signaling in PTOA, which together serve as promising therapeutic targets in pursuit of disease-modifying OA drugs.

## METHODS

### Mice

Male and female mice were housed in ventilated cages of up to five animals, given *ad libitum* access to chow food and water, on a 12 h light/dark cycle. Experimental mice were 12-16 weeks old and euthanized by CO_2_ asphyxia. C57BL/6J mice were used throughout, unless specified otherwise. Wnt-GFP reporter mice (TCF/Lef:H2B/GFP, Jax #013752) were used for flow cytometric analysis of synovial cells after joint injury and for isolation of primary synovial fibroblasts (SFs). To induce joint injury and PTOA, C57BL/6J or Wnt-GFP reporter mice were subjected to Sham (anesthesia and analgesia only) or tibial compression-based, non-invasive anterior cruciate ligament rupture (ACLR), as reported previously(21) and based on a modified protocol from Christiansen *et al*(22). Briefly, mice under inhaled isoflurane anesthesia were placed prone on a custom fixture on a materials testing system (Electroforce 3300, TA Instruments). The right knee was flexed to 100° and the paw was mounted in 30° dorsiflexion. After preloading and preconditioning, a 1.5 mm displacement was rapidly applied to the paw (10 mm/s), causing tibial subluxation and ACL rupture. Mice were then administered a single subcutaneous dose of carprofen (5 mg/kg). For R-spondin 2 delivery, mice were given intra-articular injections of recombinant mouse R-spondin 2 (500 ng, R&D Systems) or vehicle (phosphate-buffered saline, PBS) for five consecutive days using a 33G needle and microsyringe (Hamilton) in a total volume of 5 μL. Hair was removed from the hindlimbs using an electronic shaver and hair removal cream, and injections were carried out under isoflurane anesthesia. All procedures were performed according to approved IACUC protocols.

### Knee hyperalgesia testing

Prior to pain testing, mice were acclimated to handling and application of the measurement device. Knee hyperalgesia was measured in a blinded fashion using a Randall-Selitto device (IITC Life Science) modified for pressure application on a mouse knee, as described previously(21). The convex tip of the pressure applicator was applied to the medial knee joint until a vocal or physical response occurred. The average reading of triplicate applications was calculated for each limb.

### Near-infrared live imaging

To assess extracellular matrix remodeling in the joint, we employed live near-infrared imaging using the matrix metalloproteinase (MMP) activatable fluorescent probe MMPSense680 (Perkin Elmer). The day prior to imaging, hair was completely removed from the hindlimbs. Under isoflurane anesthesia, bilateral intra-articular injection of MMPsense680(48) probe (4 μL per knee) was carried out using a 33G needle and microsyringe (Hamilton). Mice were recovered from anesthesia, permitted two hours of cage activity, and then near-infrared imaging of hindlimbs was performed under anesthesia using a Pearl Impulse Imaging System (LI-COR). MMPsense signal intensity was analyzed in ImageJ using a consistent defined region of interest over the right and left knee to calculate raw integrated density value

### Histology

Limbs from mice injected with vehicle or R-spondin 2 were harvested 28 d after the first injection, fixed for 48 h in 10% neutral-buffered formalin, rinsed with water, decalcified for two weeks in 10% EDTA, then processed in paraffin. Sagittal sections (5 μM) spanning the medial joint, spaced ∼100 μM apart, were cut and stained with Safranin-O/Fast Green (SafO) and Hematoxylin. Sections were imaged at 10x magnification on a Nikon Eclipse Ni E800 microscope with a Nikon DS-Ri2 camera. Qualitative OA and synovitis grading was performed by two blinded observers according to established grading schemes(49, 50), with the addition of a fibrosis sub-score for synovitis grading. Grading criteria for PTOA and synovitis scores can be found in Supplementary Tables 1 and 2. Scores were averaged across sections within each specific compartment of interest, for each limb (minimum of three sections per limb) to obtain mean limb scores for each independent grader. Then the mean scores were averaged across graders to obtain aggregate compartment scores for each limb.

### Immunohistochemistry

Whole hindlimbs were collected from Sham or ACLR mice, fixed for 48 h in 10% neutral-buffered formalin, rinsed with water, subjected to decalcification for two weeks using 10% EDTA, and embedded in paraffin. Sagittal sections (5 μm) spanning the medial joint were cut and mounted. Epitopes were retrieved in 10 mM sodium citrate buffer (pH 6.0) for 30 min at 70°C. After permeabilization and blocking, sections were incubated overnight at 4°C with primary antibody, then with secondary antibody for 1 h at room temperature. Sections were counterstained with Hoechst 33342 (Invitrogen), then incubated in 0.2% Sudan black for 30 min and mounted with ProLong Gold (Invitrogen). Slides were imaged at 10x (at least three sections per limb) on a Lionheart FX (Biotek) for three mice per condition. For qualitative grading of R-spondin 2 staining, the following scale was used: no expression (0), <25% expression (1), 25-50% expression (2), >50% expression (3). All antibody information can be found in Supplementary Table 3.

### Immunocytochemistry

Primary SFs were plated on coverslips and fixed using 4% paraformaldehyde. Cells were permeabilized in 0.1% TBST, blocked for 1 h, then incubated overnight at 4°C in primary antibody. The following day, secondary antibody was added for 1 h at room temperature, before counterstaining with Hoechst 33342 (Invitrogen) and mounting of coverslips onto slides with ProLong Gold (Invitrogen). Images were acquired on a Lionheart FX (Biotek). All antibody information can be found in Supplementary Table 3.

### Primary synovial fibroblast isolation and culture

SFs were isolated from C57BL/6J or Wnt-GFP reporter mice at 12-16 weeks of age. Hindpaw SFs were derived from the mouse hindpaw based on previously reported methods(51). Briefly, claws and skin were removed from hindpaws, muscles and tendons excised, then longitudinal incisions were made with a scalpel along the hindpaw. Both dissected hindpaws were digested for up to 50 min at 37°C and vortexed every 5 min in a 4.5 mL volume of synovial digestion medium (DMEM with 400 μg/mL collagenase IV, 400 μg/mL liberase, 400 μg/mL DNaseI). After digestion, cells were centrifuged at 500 x *g* for 5 min then plated in DMEM containing 10% fetal bovine serum (FBS), 1x L-glutamine, and 1x antibiotic/antimycotic (Invitrogen). Knee-derived SFs were derived from mouse knee synovium as reported previously(21). Briefly, knee synovium was dissected and digested for up to 35 min while shaking at 1000 rpm at 37°C, with vortexing at 0, 15 and 30 min. Up to two synovia were digested together in a 1.5 mL volume of synovial digestion listed above. Cells were pelleted by centrifugation after digestion then plated in culture medium as listed above for hindpaw-derived SFs. Both hindpaw-derived and knee-derived SFs were used for experiments between passage 4 to 6. Prior to treatment, SFs were serum-starved overnight and treated in the absence of serum.

### Primary bone marrow-derived macrophage (BMDM) isolation and culture

BMDM were grown from whole mouse bone marrow based on previously reported protocols(21, 52). Briefly, femoral and tibial bone marrow from C57BL/6J mice aged 12-16 weeks was flushed and grown in DMEM containing 10% FBS, 1x L-glutamine, 1x antibiotic/antimycotic (Invitrogen), and 20 ng/mL recombinant M-CSF (Gibco). BMDM were lifted and seeded into experimental plates for treatment at passage 1.

### ATDC5 culture and differentiation

ATDC5 cells were kindly provided by Dr. Rhima Coleman (University of Michigan). Cells were cultured in DMEM containing 10% FBS, 1x L-glutamine and 1x antibiotic/antimycotic (Invitrogen). For chondrogenic differentiation, ATDC5 cells were grown to 90% confluency then switched to differentiation media (DMEM high glucose, 10 ng/mL TGFβ, 100 nM dexamethasone, 40 μg/mL ascorbic acid 2-phosphate, 1 mM sodium pyruvate, 40 μg/mL L-proline, and 1x ITS pre-mix containing 6.25 μg/mL insulin, 6.25 μg/mL transferrin, 6.25 μg/mL selenous acid, 5.33 μg/mL linoleic acid, 1.25 mg/mL bovine serum albumin). Cells were differentiated for 21 days, with media replaced every 3 days.

### Gene expression analysis

Transcript levels were analyzed in whole synovium, ATDC5 cells, BMDM, hindpaw-derived SFs and knee-derived SFs. Synovium were subjected to homogenization in CK28 PreCellys tubes (Bertin Technologies) containing TRIzol (Invitrogen) at 4°C for two to three cycles on the soft tissue setting (5800 rpm, 2 × 15 s, 30 s rest). Debris was then removed by centrifugation at 10,000 x *g* for 10 min before proceeding with phenol-chloroform RNA isolation, as with cell culture samples. Nucleic acid concentration and purity was determined using a Nanodrop spectrophotometer (ThermoFisher) then a standard amount of RNA subjected to reverse transcription using the High-Capacity cDNA Reverse Transcription Kit (Applied Biosystems). Quantitative real-time PCR (qPCR) was employed to assess gene expression using Power SYBR Master Mix (Applied Biosystems) on a QuantStudio5 Real-Time PCR System (Applied Biosystems). Genes of interest were normalized to expression of a stable housekeeping gene and control samples set to 1 using the 2^-ΔΔCt^ method. All primer sequences can be found in Supplementary Table 4. For calculation of the M1:M2 macrophage polarization score, based on Paige *et al*(53), we used a composite of M1 (*Il1b, Il6* and *Nos2*) and M2 (*Mrc1, Il10* and *Arg1*) gene expression.

### Protein quantification

Conditioned medium was harvested from cell cultures, centrifuged at 500 x *g* for 5 min to remove debris, then stored at -80°C. Synovial fluid was collected from mouse knees based on the protocol reported by Seifer *et al*(54). Briefly, the joint capsule was carefully opened and an alginate microsponge inserted for 30 s to recover synovial fluid. The soaked sponge was then removed and placed into a cryovial for flash freezing and storage at -80°C. Upon thawing, alginate lyase was added for incubation at 34°C for 30 min to digest the sponge. At completion, sodium citrate was added prior to vortexing.

To quantify the concentration of total protein in synovial fluid samples, a Coomassie Plus Assay Kit (based on the Bradford assay) was used in accordance with manufacturer’s instructions. This permitted normalization of target analyte concentration to total protein amount in synovial fluid, given the variability of fluid recovery from mouse to mouse. To measure the concentration of R-spondin 2 in synovial fluid or cell culture conditioned medium, a mouse R-spondin 2 ELISA kit (CusaBio) was used according to manufacturer’s instructions.

### Flow cytometry

Multicolor flow cytometry was used to assess synovial cell types based on expression of surface-based or intracellular proteins, to measure cell viability, or to assess endogenous reporter signal. Synovial tissue was dissected and digested as described above, resulting in a single cell suspension. For viability testing, cells were either pre-stained with a fixable viability dye (eFluor660, eBioscience) or post-stained with a nuclear viability dye (TOPRO3, Invitrogen). Cold FACS buffer (PBS containing 2% FBS, 1 mM EDTA) was used for all staining and preparation steps, and for running samples on the cytometer. For all experiments, non-specific binding was blocked using TruStain FcX (Biolegend). For intracellular detection of R-spondin 2, cells were fixed and permeabilized using the Cyto-Fast Fix-Perm Buffer Set (Biolegend), and an anti-R-spondin 2 antibody (ProteinTech) or Rabbit IgG Isotype Control (Abcam) were used with a secondary anti-rabbit BV421. CD31-PECy7 (Biolegend) and CD45-BV650 (Biolegend) were used for surface staining to identify major synovial cell types, and the Wnt-GFP fluorescent reporter was used to assess canonical Wnt signaling activity based on TCF/Lef transcriptional activity. Antibodies and their conjugates are listed in detail in Supplementary Table 3. In addition to fully stained samples, all experiments included an unstained cells control, single-stained controls comprised of cells or UltraComp eBeads (Invitrogen), and fluorescence-minus-one (FMO) or isotype controls to determine population gating. Compensation and acquisition were performed using a BD LSRFortessa cytometer with FACSDiva software (BD Biosciences), then data analysis was performed using FlowJo v10 (TreeStar/BD Biosciences).

### Single-cell RNA-sequencing (scRNA-seq)

For scRNA-seq, two biological replicates of the following conditions were used: Sham (no injury, anesthesia/analgesic only), 7d ACLR, and 28d ACLR, for a total of six samples. Each biological replicate was comprised of a synovium from one male and from one female mouse. Synovia were dissected and digested as described above, to yield a single cell suspension at >90% viability. Single cells were immediately submitted to the University of Michigan Advanced Genomics Core for loading onto the 10x Genomics pipeline (Chromium Next GEM Single Cell 3’ Kit v3.1) for barcoding and library preparation. Pooled libraries were then submitted to BGI for paired-end (100bp+100bp) sequencing on the DNBseq G400 system (MGI Tech), generating ∼350 million reads per sample (>50 000 reads per cell). Raw data were aligned to the reference mouse transcriptome using STAR(55) and underwent pre-processing in Cell Ranger (10x Genomics).

### Quality control, filtering, clustering, annotation, and analysis of scRNA-seq data

Pre-processed aligned data were read from Cell Ranger into Seurat (R, v4.1.0). Cells with less than 100 detected genes and genes expressed in less than 10 cells were excluded from Seurat object generation. Next, low-quality cells were filtered out based on number of expressed features (i.e. genes), with remaining cells having 200 < *nFeature* < 6000. Cells in which > 5% of all genes were mitochondrial-derived (percent.mt > 0.05) were also excluded. Canonical correlation analysis, which performs anchor-based integration(56), was used to integrate data across conditions (Sham, 7d ACLR, 28d ACLR) using the top 3000 variable features. In tandem, we performed Spearman correlation analysis of average expression for all genes between integrated biological replicates of each condition. *ScaleData()* was used to perform linear transformation and to regress out unwanted variation due to cell cycle and mitochondrial gene expression. Linear dimensionality reduction was performed using principal component analysis, and elbow plots and principal component heatmaps were used to determine the number of dimensions used in subsequent nonlinear dimensionality reduction via Uniform Manifold Approximation and Projection (UMAP). Dimensions 1-25 were used in the object representing all conditions, all biological replicates, and all cells. Unsupervised clustering was performed using *FindNeighbors* (dims = 1:25) and *FindClusters* (resolution = 0.3) to derive distinct cell clusters.

Cluster markers were identified using *FindAllMarkers*. Genes expressed in greater than 80% of cells in the cluster of interest and in less than 20% of cells in all other cells were focused on, as were genes with positive differential expression relative to other clusters, as opposed to negative markers. To identify and functionally annotate cell clusters, the genes outputted by *FindAllMarkers* for each cluster were submitted to Gene Ontology (GO) pathway analysis (GO: Biological Process) using PantherDB(57, 58). We further utilized Cluster Identity Predictor (CIPR)(59) and published synovial scRNA-seq datasets at the Single Cell Portal(4) to compare gene marker profiles to known cell annotations. Violin plots and gene feature plots were generated using the Seurat functions *VlnPlot()* and *FeaturePlot()*, respectively. Bubble plots of GO terms were made using ggplot2(60). Global gene expression differences between conditions or groups were assessed by identifying the top 25 differentially expressed genes (DEGs) between conditions with most positive and most negative log2FC using a pseudo-bulk analysis approach and DESeq2(61). Read counts of genes were then subset by sample, and then averaged for all cells across each sample. Global z-scores of average read count for each gene were computed and plotted using *pheatmap*.

To identify fibroblast subsets, all fibroblast-like cells (not including pericytes) were subset from the “all synovial cells” Seurat objects within each condition and re-clustered (dims = 1:25, resolution = 0.13). Cluster markers and functional annotation were performed as described above. To identify conserved functions across all fibroblast subsets, the *gsfisher* R package was used to test for GO overrepresentation, and the gene universe (i.e. background list of expressed genes) was defined as genes expressed in 25% of cells in the cluster of interest or 25% of all remaining cells. To identify unique functions across fibroblast subsets, the statistically significant (*Q* < 0.05, where *Q* is the *P* value adjusted for false discovery rate) genes and their Log2FC from *FindAllMarkers()* were submitted to GO (Biological Processes) or Reactome analysis via PantherDB. Pathway terms were manually curated and grouped into categories of conserved themes based on the following criteria: *Q* < 0.05 (expressed as log_10_(FDR)), only found in a single cluster, rich ratio (higher number of DEGs in proportion to the total genes in term), and pathway term similarity. Fibroblast subsets were identified, defined, and named based on these categories and known, functionally relevant cluster markers.

### Intercellular communication analysis

The CellChat R package (CellChat 1.1.3)(27) was used to infer and quantify cell-cell communication networks. In the analysis of all synovial cells, macrophages and dendritic cells were combined into a single “myeloid” group, and vascular endothelial cells and lymphatic endothelial cells were combined into a single “endothelial” group to focus on major cell group signaling. Consistent with recommendations by the CellChat toolbox to curate expanded ligand-receptor sets for focused analysis of signaling axes of interest, an “Rspo” pathway of known ligand-receptor interactions between R-spondins 1-4 and Lgr receptors 4-6 was added to the pathway interaction database. Over-expressed genes and ligand-receptor interactions were identified with default *identifyOverExpressedGenes()* and *identifyOverExpressedInteractions()* functions. Data were projected against the provided PPI.mouse dataset to reduce dropout effects of signaling genes. Probabilities of cell-cell communications were inferred with *computeCommunProb()*. The method for calculating average gene expression per cell group was *triMean* for all synovial cells, and *truncatedMean* with 10% trim for SFs. River plots were generated with pattern numbers at which cophenetic and silhouette measure scores decreased simultaneously, and conservative cutoff values at which all patterns were present. Circle plots and contribution plots for pathways of interest were constructed using the *netVisual_aggregate()* and *netAnalysis_contribution()* functions.

### Trajectory and Transcription Factor Binding Motif Analyses

Cellular trajectory analysis was performed using Monocle3 (v1.0.0)(62) in R. First, the Seurat object representing all fibroblasts across all conditions was imported into Monocle3 using the Seurat-Monocle wrapper. Data was preprocessed with the same number of dimensions as the Seurat-based analysis, and nonlinear dimensionality reduction was performed using UMAP. Trajectories were reconstructed using the *learn_graph()* function, and cells were ordered along pseudotime using the *order_cells()* function. The reconstructed trajectory identified a terminal node in the Dpp4+ cluster, which was selected as the trajectory root. Next, the trajectory branch originating in the Dpp4+ cluster and terminating in the Prg4^hi^ cluster was subset for focused pseudotime-dependent analyses of Prg4^hi^ lining fibroblasts. Unsupervised preprocessing, dimensionality reduction, and clustering of this trajectory was performed, and *learn_graph()* identified a trajectory from Dpp4+ cells to Prg4^hi^ cells, via αSMA+ cells. The trajectory node in the most proximal aspect of the Dpp4+ cluster was again selected as a root, and pseudotime was calculated. To find genes that vary as a function of pseudotime along a trajectory, gene read count data was fitted to a quadratic spline function, similar to the *plot_genes_in_pseudotime()* function in Monocle3. Genes with quadratic r^2^ > 0.15 and with trajectory-wide average read count values > 0.1 (to exclude noise) were then analyzed for co-regulation using the *find_gene_modules()* function, which groups genes into modules depending on their variance patterns along pseudotime using Louvain community analysis. Modules with high degree of composite expression in the Prg4^hi^ cluster (Module 2 in Figure S8I), representing genes that either increase or decrease with similar variance patterns indicative of co-regulation, were selected for subsequent transcription factor (TF) binding analysis to identify the TFs regulating lineage specification of Prg4^hi^ lining cells. These genes were then input to the RcisTarget(63) R package for DNA binding motif analysis. *mm9-tss-centered-10kb-7species.mc9nr.feather* was applied as a database with motifs using the “directAnnotation” parameter. SOX5 was predicted as a key TF enriched in the Prg4^hi^ lining cluster that regulates *Rspo2* and other co-regulated genes along the differentiation trajectory. TF gene regulatory network for SOX5 was then visualized using Cytoscape (v3.91)(64).

### Sample sizes and statistical analyses

Microsoft Excel was used for tabulation of raw data. GraphPad Prism 9 was used for statistical analyses and generation of figures. All data are presented as mean ± SEM and *P*<0.05 was considered statistically significant. Shapiro-Wilk tests were used to assess normal distribution. For two-group comparisons of normally distributed data, parametric two-tailed student’s t-tests were used, in a paired or unpaired manner dependent upon the experimental design. For comparisons of data across three or more independent groups/conditions, one-way or two-way ANOVA with post-hoc Tukey testing were used for normally distributed data. For scRNA-seq analyses, we used default statistical tests built into R packages Seurat, Monocle, RcisTarget, and CellChat, that employ false discovery rate testing to account for multiple comparisons in large datasets.

## Acknowledgements

We would like to acknowledge the assistance of the University of Michigan Advanced Genomics Core, Flow Cytometry Core, Orthopaedic Research Laboratories Histology Core, and Unit for Laboratory Animal Medicine. We appreciate the intellectual input and critical feedback from Drs. Ormond MacDougald and Jan Stegemann. BioRender.com was used to generate cartoons and schematics. Data in this manuscript were presented at the Orthopaedic Research Society’s Annual Meetings in 2021 and 2022.

## Contributors

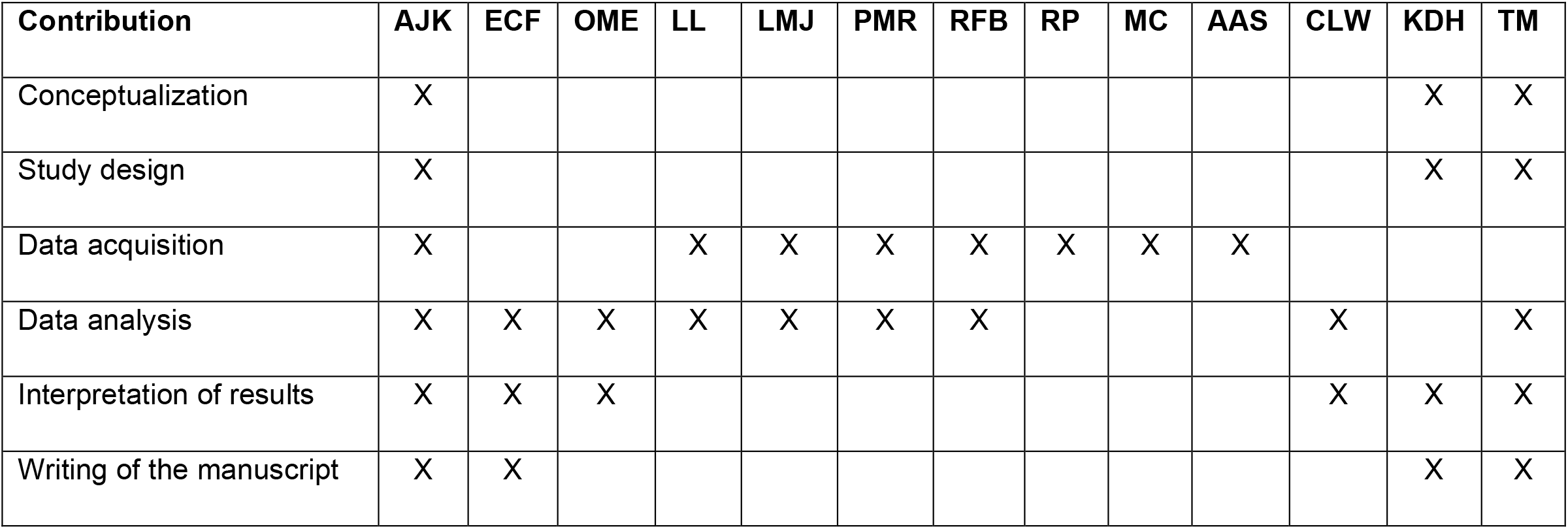

## Ethics approval

All animal experiments were performed in accordance with approved IACUC protocols.

## Data availability statement

All sequencing data will be uploaded to the NCBI Gene Expression Omnibus (GEO) and made public before the manuscript is in press.

