## Supplementary Material for "Synovial fibroblasts assume distinct functional identities and secrete R-spondin 2 to drive osteoarthritis"

**SUPPLEMENTAL MATERIAL**

**SUPPLEMENTARY FIGURES**

**Figure S1. Viability testing and quality control of scRNA-seq data.** (A) Flow cytometry analysis of synovial cells to assess major cell populations and viability. Small debris were first excluded (1) then single cells gated on (2,3). Dead cell exclusion (4, including mean percentage of live cells) was performed, and the resulting live single synovial cells analyzed for expression of CD31 (5, endothelial cells), CD45 (6, hematopoietic cells), or lack of expression of CD31 and CD45 (7, double negative cells including predominantly SFs). (B) Schematic illustrating the experimental design of the scRNA-seq experiment. (C) Quality control and filtering steps prior to analysis of scRNA-seq data. Corresponding plots are included for each step. (D) Cluster-by-cluster Spearman correlation analysis of biological replicates for each condition. Color bar represents the Spearman correlation co-efficient of the average expression for all genes from each replicate (ρ). FSC-A: forward scatter-area; SSC-A: side scatter-area; FSC-W: forward scatter-width; SSC-W: side scatter-width; FSC-H: forward scatter-height; SSC-H: side scatter-height.

**Figure S2. Markers and annotation of all synovial cell types.** (A) Heatmap showing the top 5 genes per cluster as determined by FindAllMarkers in Seurat, corresponding to clusters in UMAP plots from Figure 1A, ranked by log2FC. (B) UMAP plot showing all synovial cells from all conditions integrated by canonical correlation analysis. Major cell groups have dashed outlines and cell type annotations are shown on the right.

**Figure S3. Outgoing and incoming communication patterns for all synovial cells.** (A) Line plots for determining the number of outgoing communication patterns in CellChat analysis. (B) Heatmap of outgoing communication patterns for each cell type. (C) Heatmap showing the contribution of each signaling pathway to each outgoing communication pattern. (A-C) correspond to the river plot in Figure 1F. (D) Line plots for determining the number of incoming communication patterns. (E) Heatmap of incoming communication patterns for each cell type. (F) Heatmap showing the contribution of each signaling pathway to each incoming communication pattern. (G) River plot showing CellChat analysis of incoming signaling patterns to major synovial cell types and the pathways comprising each pattern, with contributions scores of each signaling pathway shown on the right. (D-F) correspond to the river plot in Figure S3G.

**Figure S4. Markers and annotation of synovial fibroblast subsets.** (A) Heatmap showing the top 5 genes per cluster as determined by FindAllMarkers in Seurat, ranked by log2FC. Corresponding to clusters in UMAP plots from Figure 2A-B. (B) UMAP plot of SFs showing cluster borders with colored, dashed outlines. Duplicated elsewhere throughout figures. (C) Feature plots of additional SF subset marker genes for each cluster determined by FindAllMarkers analysis in Seurat. Color scales represent relative expression for each gene individually. OCP: osteochondral progenitor.

**Figure S5. Functions and communication patterns of synovial fibroblast subsets.** (A) Pathway analysis using a GeneUniverse approach (with GO Biological Processes terms) to assess conserved functions across SF clusters. (B) Reactome pathway analysis showing unique functional terms for each cluster. (C) Line plots for determining the number of outgoing communication patterns. (D) Heatmap of outgoing communication patterns for each SF cluster. (E) Heatmap showing the contribution of each signaling pathway to each outgoing communication pattern in Figure 2I. (C-E) correspond to the river plot in Figure 2I. (F) Line plots for determining the number of incoming communication patterns. (G) Heatmap of incoming communication patterns for each SF cluster. (H) Heatmap showing the contribution of each signaling pathway to each incoming communication pattern for each SF cluster. (I) River plot showing CellChat analysis of incoming signaling patterns to SF clusters and the pathways comprising each pattern, with contribution score shown on the right. (F-H) correspond to the river plot in Figure S5I. n_fg: number of foreground genes found in a GO term, FDR: false discovery rate.

**Figure S6. Wnt signaling is induced in synovium after joint injury.** (A) Relative contribution of Wnt pathway ligand-receptor pairs for the circle plot of all synovial cells in Figure 3A. (B) Relative contribution of Wnt pathway ligand-receptor pairs for the circle plot of SFs in Figure 3B. (C-D) Pseudobulk heatmaps of significantly differentially expressed genes (DEGs) from the GO term cell-cell signaling by Wnt (GO:0198738) in (C) all synovial cells or (D) SFs across conditions. Color scale (z-score) of expression for C and D is located above heatmaps.

**Figure S7. Flow cytometric analysis of synovium from injured Wnt-GFP reporter mice.** (A) Strategy for gating live Wnt-GFP+ synovial cells by flow cytometry using a fluorescence-minus-one (FMO) control lacking endogenous Wnt-GFP (left) and a fully stained sample from a Wnt-GFP reporter mouse (right). TOPRO3 was used as a dead cell exclusion dye. (B) Wnt-GFP+ cells as a percentage of all synovial cells in contralateral (contra) and 7d ACLR synovium (n=4 mice). (C) Wnt-GFP+ CD31- CD45- cells as a percentage of all CD31- CD45- cells in contra and 7d ACLR synovium (n=4 mice). ns: not significant. Error bars are mean ± SEM.

**Figure S8. R-spondin 2 is activated in synovium following joint injury.** (A) mRNA expression of *Rspo2* in 3d or 14d ACLR and contralateral synovium (n=6-8 mice per condition). *Rspo2* levels were normalized to *Atp5b* and the 3d contra condition set to 1. (B) Total number of CD31- CD45- cells expressing RSPO2 in contra, 7d or 28d ACLR synovium (n=3 mice per condition). (C) Relative expression of *Rspo* family member genes in SFs across conditions. (D) Feature plots of *Rspo1-4* in SFs with corresponding cluster outlines (right). Color scales are not equivalent between plots. (E) mRNA expression of *Rspo2* in cultured BMDM and knee-derived SFs, normalized to *Atp5b* and with BMDM set to 1 (n=3 biological replicates). (F) Immunofluorescent staining of RSPO2 in cultured knee-derived SFs. A secondary antibody (Ab) only control is shown (left) and nuclei were counterstained with DAPI. (G) Relative contribution of Rspo pathway ligand-receptor pairs for the circle plot of SFs in Figure 4H. (H) Pseudotime plot for the Dpp4+ to $Prg4^{hi}$ lining SF trajectory in Figure 4I, calculated using Monocle3. (I) Heatmap of co-regulated gene modules for the Dpp4+ to $Prg4^{hi}$ lining trajectory. For (A), unpaired two-tailed student's t-tests were used to compare expression in contra versus ACLR synovium at each time point, where ***P*<0.01 compared to contralateral. For (B), two-way ANOVA with multiple comparisons and Tukey's post-hoc testing was used, where **P*<0.05, ****P*<0.001 compared to contralateral. For (E), an unpaired two-tailed student's t-test was used where **P*<0.05. For A, B and E, error bars are mean ± SEM.

**Figure S9. R-spondin 2 orchestrates crosstalk between synovial fibroblasts and macrophages.** (A) Relative expression of *Lgr* family member genes in SFs across conditions, scaled for each gene individually, from scRNA-seq. (B) qPCR analysis of relative expression of *Lgr4-6* in hindpaw-derived SFs (left, n=7 biological replicates) and knee-derived SFs (right, n=10 biological replicates), with levels normalized to *Atp5b* and *Lgr4* set to 1. (C) Knee-derived SFs were treated for 24 h with vehicle, RSPO2 (200 ng/mL), Mianserin (20 µM) or RSPO2+Mianserin (n=3-10 biological replicates). Expression of *Axin2*, *Lef1* and *Rspo2* was measured by qPCR and normalized to *Atp5b* levels, with vehicle-treated samples set to 1. (D) Feature plots of *Lgr4-6* mapped onto all synovial cells showing distribution of expression. Major cell types are indicated. Color scales for expression of each gene are not equivalent. For B and C, one-way ANOVA with multiple comparisons and Tukey's post hoc testing was performed, where **P*<0.05, ***P*<0.01, ****P*<0.001. Error bars are mean ± SEM.

**Figure S10. R-spondin 2 orchestrates crosstalk between synovial fibroblasts and chondrocytes.** (A) Relative expression of *Lgr4-6* in ATDC5 cells following 21 d of chondrogenic differentiation (n=8), with levels normalized to *Gapdh* and *Lgr4* set to 1. (B) Conditioned media from SFs treated for 24 h with vehicle (veh) or RSPO2 (200 ng/mL) was given to differentiated ATDC5 cells for 24 h (n=3). Expression of osteogenic genes *Bsp* and *Ocn* was measured by qPCR and normalized to *Gapdh* levels, with veh-treated samples set to 1. (C) ATDC5 cells were differentiated for 21 d with veh or RSPO2 (200 ng/mL) (n=6). Expression of *Acan*, *Col2a1*, *Col10a1*, *Runx2*, *Ocn*, *Osx* and *Axin2* was measured by qPCR and normalized to *Gapdh* levels, with veh-treated samples set to 1. (D) ATDC5 cells were differentiated for 21 d then treated for 24 h with veh or RSPO2 (200 ng/mL) (n=8). Expression of *Acan*, *Col2a1*, *Col10a1*, *Ocn*, *Bsp*, *Axin2* and *Lef1* was measured by qPCR and normalized to *Gapdh* levels, with veh-treated samples set to 1. For (A), one-way ANOVA with multiple comparisons and Tukey's post hoc testing was performed, where *$P<0.05$, **$P<0.01$, ***$P<0.001$. For B-D, unpaired two-tailed student's t-tests were used, where *$P<0.05$, **$P<0.01$, ***$P<0.001$. Error bars are mean ± SEM.

**SUPPLEMENTARY TABLES**

**Supplementary Table 1. PTOA severity scoring guidelines, as published previously(1, 2)**

| **Feature** | **0** | **1** | **2** | **3** | **4** | **5** | **6** | **7** |
|---|---|---|---|---|---|---|---|---|
| **Structural Damage (0-7)** | Normal Cartilage | Roughened surface with small fibrillations | Fibrillations immediately below superficial layer or some loss of laminal surface | Horizontal cracks or separations between calcified and noncalcified cartilage | Mild loss of non-calcified cartilage (<10% surface area) | Moderate loss of non-calcified cartilage (10-50% surface area) | Severe loss of non-calcified cartilage (>50% surface area) | Erosion of cartilage to subchondral bone (any percent of surface area) |
| **Proteoglycan Loss (0-3)** | Normal cartilage | Decreased but not complete loss of toluidine blue staining in non-calcified area | Focal loss of toluidine blue staining in non-calcified cartilage (<30% surface area) | Diffuse loss of toluidine blue staining in non-calcified cartilage (>30% surface area) | | | | |
| **Chondrocyte Hypertrophy (0-1)** | None | Enlarged chondrocyte lacunae with lack of toluidine blue stain around a collapsed cell | | | | | | |
| **Osteophyte Size (0-3)** | None | Small- same thickness as adjacent cartilage | Medium- 1 to 3 times thick as adjacent cartilage | Large- >3 times thicker than adjacent cartilage | | | | |
| **Osteophyte Maturity (0-3)** | None | Predominately cartilage | Mixed cartilage and bone with vascular invasion | Predominately bone | | | | |
| **SCB Thickening (0-3)** | Normal cartilage | Mild thickening, <50% increase | Moderate thickening, 50-100% increase | Severe thickening, >100% increase | | | | |

**Supplementary Table 2. Synovitis scoring guidelines, as published previously(2, 3), with the addition of a fibrosis criterion**

| Feature | Description | 0 | 1 | 2 | 3 |
|---|---|---|---|---|---|
| Pannus | Defined as fibrous tissue/synovium/ inflammatory cell outgrowth spreading over the surface of the bone and/or cartilage at the joint margins. | No pannus | Mild: Pannus has migrated on bone but not encroaching on cartilage. | Moderate: Pannus has migrated < 1x cartilage depth. | Severe: Pannus has migrated > 1x cartilage depth. |
| Bone erosion | Score the margin of the femur at or adjacent to the joint capsule attachment (between joint capsule attachment and cartilage margin – above the junction of the growth plate with the femoral cortex). | No Cortical Bone Erosion | Partial thickness loss of cortical bone only. | Focal complete loss of cortical bone – communication with marrow cavity at one small "vascular" communication site. | Widespread complete loss of cortical bone – communication with marrow cavity at multiple sites or broad area loss of cortical bone. |
| Synovial lining hyperplasia | Score this superior to the meniscal remnant. Do not score the cells actually attached to the tibia or femur or cells on the surface of the meniscus itself and avoid peri-meniscal plica. Score the maximum hyperplasia seen anywhere along this area. | 1 cell thick | Mild: 2-3 cells thick | Moderate: 4-5 cells thick | Severe: > 6 cells thick |
| Sub-synovial inflammation | Score this superior to the meniscal remnant. The infiltration of inflammatory cells (neutrophils, macrophages and/or lymphocytes) is evaluated. | No inflammatory cells | Occasional scattered inflammatory cells – or perivascular | Focal areas of dense sub-synovial WBC infiltrate – but still predominantly normal sub-synovial areolar connective tissue present. | Widespread dense sub-synovial WBC infiltrate – markedly reduced or little/no normal areolar connective tissue evident or lymphoid follicle formation. |
| Synovial fibrosis | Score this as a function of synovial area exhibiting pathological fibrosis, characterized by blue staining of collagen (from FastGreen) and reduction of areolar-like adipocytes in fat pad. | No fibrosis | Dispersed fibrosis (<10% of synovial area) | Moderate fibrosis (10-30% of synovial area) | Severe fibrosis (>30% of synovial area) |
| Synovial exudate | Infiltration of inflammatory cells (neutrophils, macrophages and/or lymphocytes) in the synovial cavity. | No inflammatory cells or fibrin in the synovial cavity. | Inflammatory cells and/or fibrin clot in the synovial cavity – may be restricted to recesses. | | |

**Supplementary Table 3. Antibodies**

| Antibody name | Raised in | Manufacturer | RRID | Application (dilution) |
|---|---|---|---|---|
| BV650 anti-mouse CD45 Clone 30-F11 | Rat | Biolegend 103151 | AB_2565884 | Flow cytometry, 1:400 |
| PECy7 anti-mouse CD31 Clone 390 | Rat | Biolegend 102417 | AB_830756 | Flow cytometry, 1:200 |
| TruStain FcX PLUS (anti-mouse CD16/32) Clone S17011E | Rat | Biolegend 156604 | AB_2783138 | Flow cytometry, 1:1000 |
| Anti-mouse R-spondin 2 Polyclonal | Rabbit | ProteinTech 17781-1-AP | AB_2269700 | Flow cytometry, 2 µg/sample; IHC/ICC, 1:250 |
| Rabbit IgG Isotype Control Polyclonal | Rabbit | Abcam ab171870 | AB_2687657 | Flow cytometry, 2 µg/sample |
| BV421 anti-rabbit IgG secondary Polyclonal | Donkey | Biolegend 406410 | AB_10897810 | Flow cytometry, 1:100 |
| AlexaFluor488 anti-rabbit Polyclonal | Goat | Invitrogen A-11008 | AB_143165 | IHC/ICC, 1:500 |

**Supplementary Table 4. Primers**

| Gene name | 5' – 3' sequence | Organism |
|---|---|---|
| *Acan* Fwd | CCTGCTACTTCATCGACCCC | *Mus musculus* |
| *Acan* Rev | AGATGCTGTTGACTCGAACCT | *Mus musculus* |
| *Arg1* Fwd | ATCGTGTACATTGGCTTGCG | *Mus musculus* |
| *Arg1* Rev | GGCCTTTTCTTCCTTCCCAG | *Mus musculus* |
| *Atp5b* Fwd | CTGGATTCAGGGGCACCAAT | *Mus musculus* |
| *Atp5b* Rev | GCACCTCCAAAGAGTCCGAT | *Mus musculus* |
| *Axin2* Fwd | GCGCTTTGATAAGGTCCTGG | *Mus musculus* |
| *Axin2* Rev | TCATGTGAGCCTCCTCTCTTT | *Mus musculus* |
| *Bsp/Ibsp* Fwd | TTCGTTTGAAGTCTCCTCTTCC | *Mus musculus* |
| *Bsp/Ibsp* Rev | CTCCTCTGAAACGGTTTCCA | *Mus musculus* |
| *Col2a1* Fwd | ACTTGCCAAGACCTGAAACTCTG | *Mus musculus* |
| *Col2a1* Rev | AAACTTTCATGGCGTCCAAGG | *Mus musculus* |
| *Col10a1* Fwd | TCTCCCAGCACCAGAATCTATC | *Mus musculus* |
| *Col10a1* Rev | CTTTATGCCTGTGGGCGTTT | *Mus musculus* |
| *Gapdh* Fwd | GCCTCTCTTGCTCAGTGTCC | *Mus musculus* |
| *Gapdh* Rev | CTCCCACTCTTCCACCTTCG | *Mus musculus* |
| *Il10* Fwd | GCGCTGTCATCGATTTCTCC | *Mus musculus* |
| *Il10* Rev | ATGGCCTTGTAGACACCTTGG | *Mus musculus* |
| *Il1b* Fwd | TGCCACCTTTTGACAGTGATG | *Mus musculus* |
| *Il1b* Rev | AAGGTCCACGGGAAAGACAC | *Mus musculus* |
| *Il6* Fwd | GCCTTCTTGGGACTGATGCT | *Mus musculus* |
| *Il6* Rev | TGCCATTGCACAACTCTTTTCT | *Mus musculus* |
| *Lef1* Fwd | AAGAAATGAGAGCGAATGTCGT | *Mus musculus* |
| *Lef1* Rev | TTCTGGGACCTGTACCTGAAGT | *Mus musculus* |
| *Lgr4* Fwd | GCTGCGGACTCTGGACTTAT | *Mus musculus* |
| *Lgr4* Rev | TGTCCCAAGCTTCGCAAAAG | *Mus musculus* |
| *Lgr5* Fwd | ATCAGGTCAATACCGGAGCG | *Mus musculus* |
| *Lgr5* Rev | GGCACCATTCAAAGTCAGTGT | *Mus musculus* |
| *Lgr6* Fwd | GGCATCATGCTGTCCGC | *Mus musculus* |
| *Lgr6* Rev | GGTTGTTCATACTGAGGTCTAGGT | *Mus musculus* |
| *Mrc1/Cd206* Fwd | GGCTGATTACGAGCAGTGGA | *Mus musculus* |
| *Mrc1/Cd206* Rev | ATGCCAGGGTCACCTTTCAG | *Mus musculus* |
| *Nos2* Fwd | AGAGAACGGAGAACGTTGGA | *Mus musculus* |
| *Nos2* Rev | GGATTCTGGAACATTCTGTGCTG | *Mus musculus* |
| *Ocn/Bglap* Fwd | CAGACACCATGAGGACCATCTT | *Mus musculus* |
| *Ocn/Bglap* Rev | GATAGCTCGTCACAAGCAGG | *Mus musculus* |
| *Osx/Sp7* Fwd | TTCCCCAGGGTTGTTGAGTC | *Mus musculus* |
| *Osx/Sp7* Rev | ATGGCGTCCTCTCTGCTTGA | *Mus musculus* |
| *Rspo2* Fwd | ACGTGTAGCAGAAACAACCG | *Mus musculus* |
| *Rspo2* Rev | GCTTCCGCCTCTTCTTCTTG | *Mus musculus* |
| *Runx2* Fwd | CCCAGCCACCTTTACCTACA | *Mus musculus* |
| *Runx2* Rev | TATGGAGTGCTGCTGGTCTG | *Mus musculus* |
| *Tnf* Fwd | ATGGCCTCCCTCTCATCAGT | *Mus musculus* |
| *Tnf* Rev | TGGTTTGCTACGACGTGGG | *Mus musculus* |
