## Supplementary Figure 1 for "Synovial fibroblasts assume distinct functional identities and secrete R-spondin 2 to drive osteoarthritis"

### Fig S1 - Viability testing and quality control of scRNA-seq data

**A** Synovial cell viability and gating

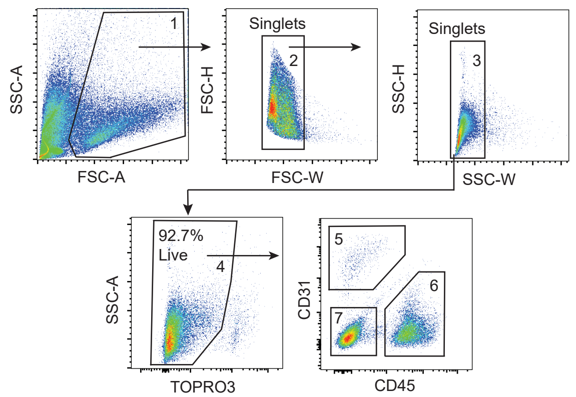


**B** scRNA-seq experimental design

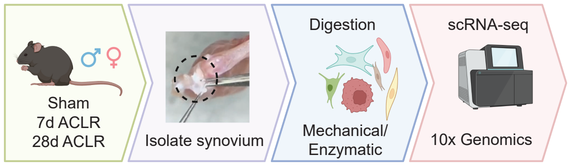


**C** scRNA-seq quality control and filtering

| Step | Number of cells | % cells remaining |
|---|---|---|
| **Step 0:** Starting number of cells in pre-processed data | 28,277 | 100 |
| **Step 1:** Load pre-processed data into Seurat min.features = 100 | 28,023 | 99.1 |
| **Step 2:** Removal of cells with nFeatures < 200 | 27,743 | 98.11 |
| **Step 3:** Removal of cells with over 5% of genes mitochondrial-derived percent.mt > 0.05 | 20,664 | 73.07 |
| **Step 4:** Removal of cells with nFeatures > 6000 | 20,422 | 72.22 |

**D** Biological replicate correlation

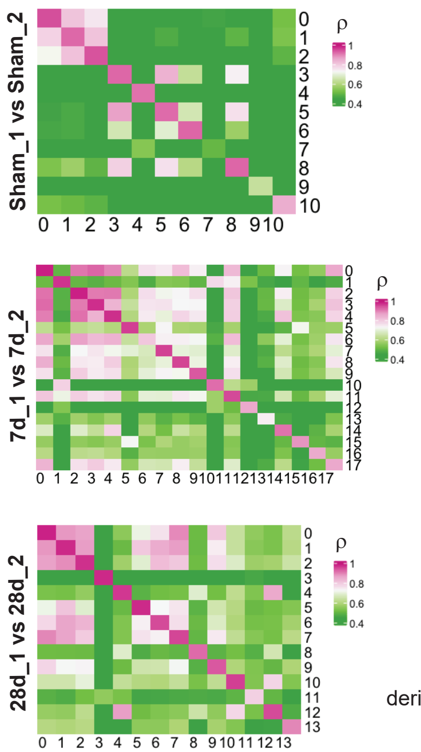


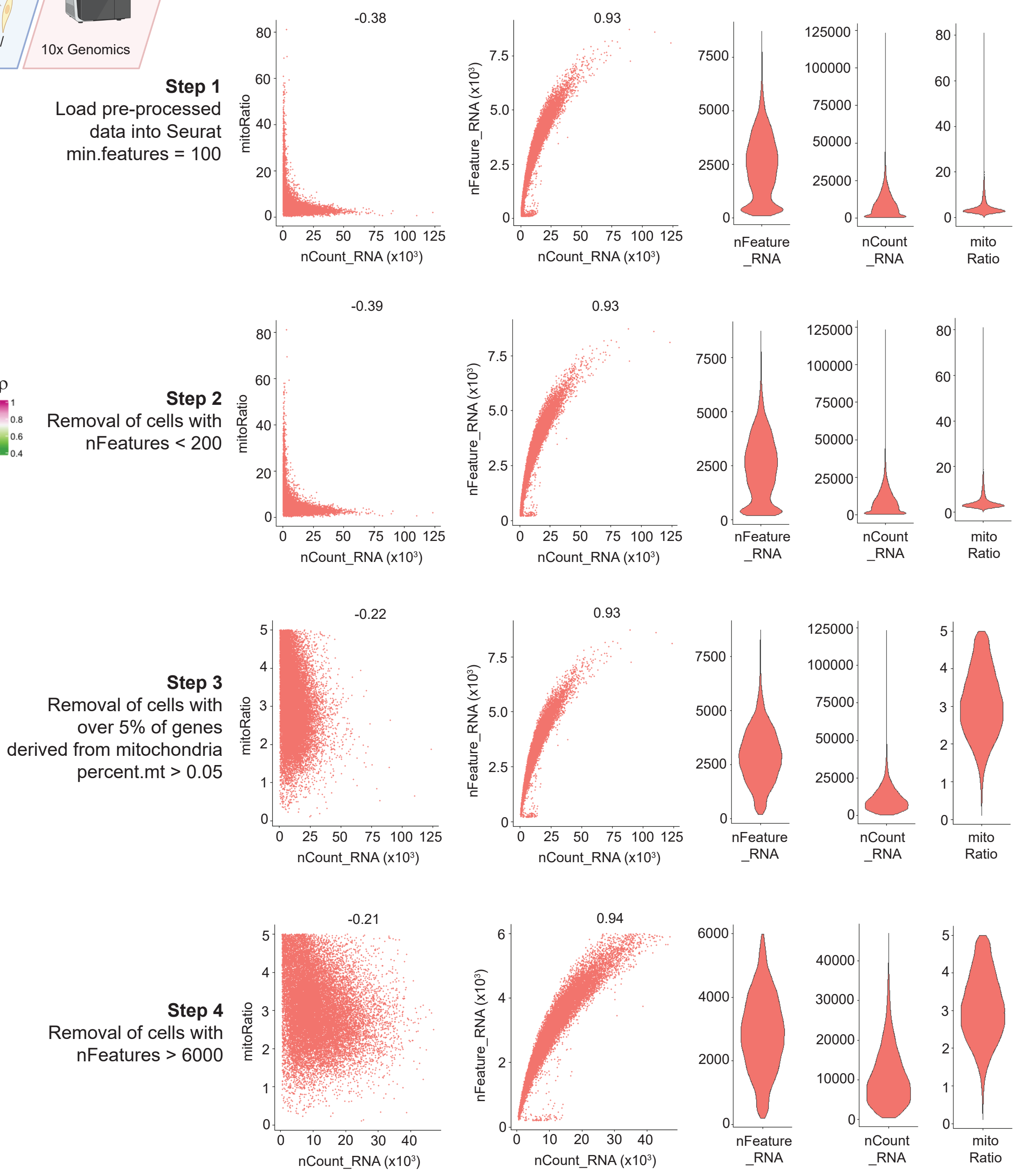
