## Supplementary Figure 2 for "Synovial fibroblasts assume distinct functional identities and secrete R-spondin 2 to drive osteoarthritis"

### Fig S2 - Markers and annotation of all synovial cell types

**A**

**All synovial cells: top 5 genes per cluster**

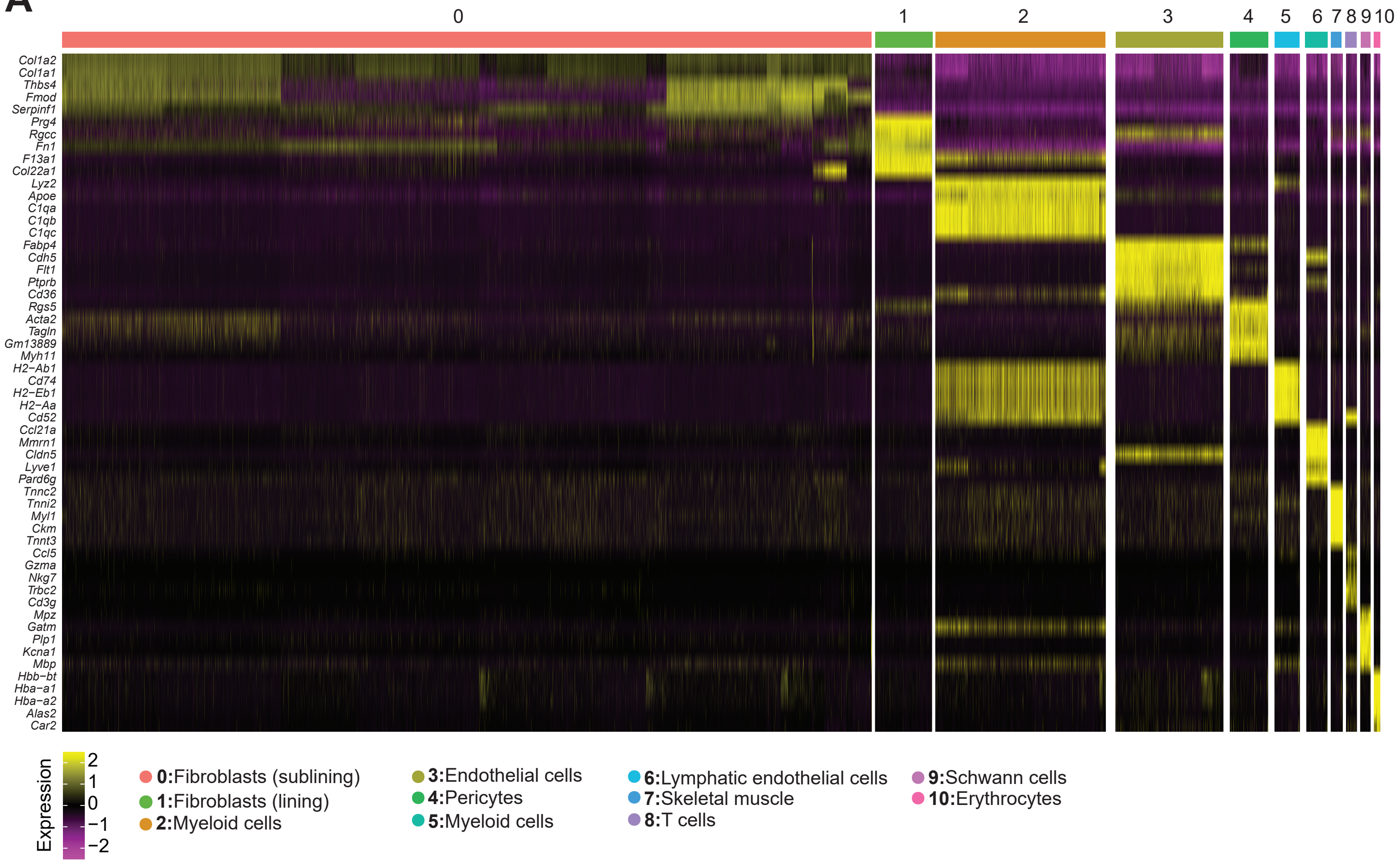


**B**

**All synovial cells, all conditions integrated**

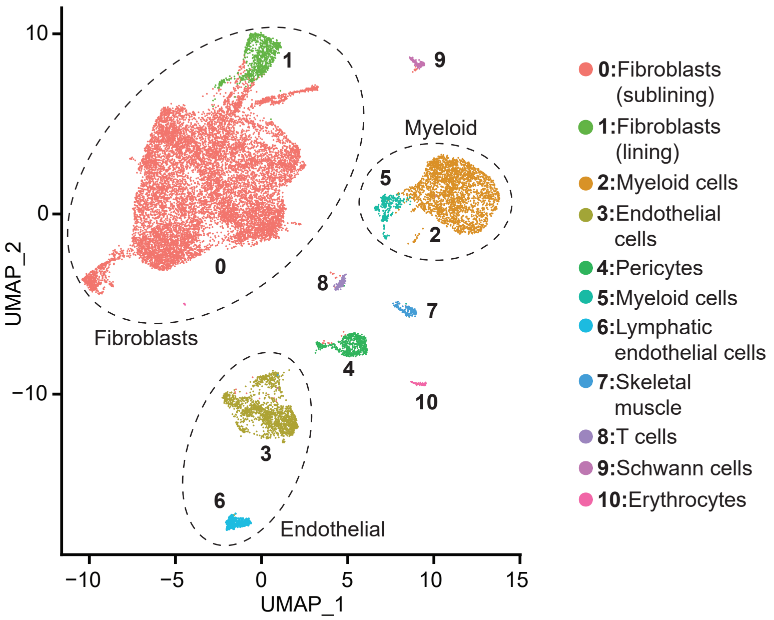
