## Supplementary figures and images for "Synovial fibroblasts assume distinct functional identities and secrete R-spondin 2 to drive osteoarthritis"

### Supplementary Figure 3

#### Fig S3 - Outgoing and incoming communication patterns for all synovial cells

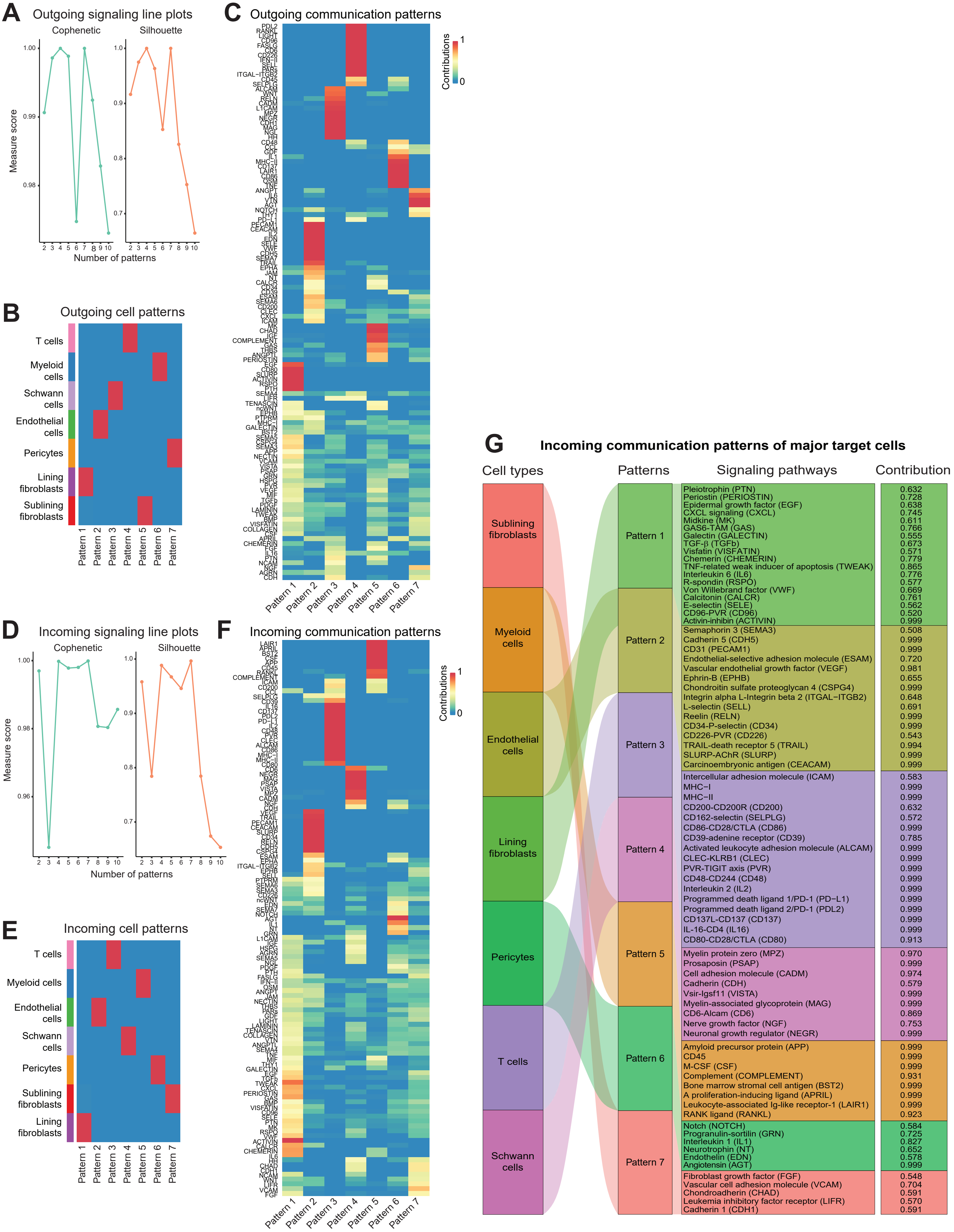

### Supplementary Figure 4

#### Fig S4. Markers and annotation of synovial fibroblast subsets

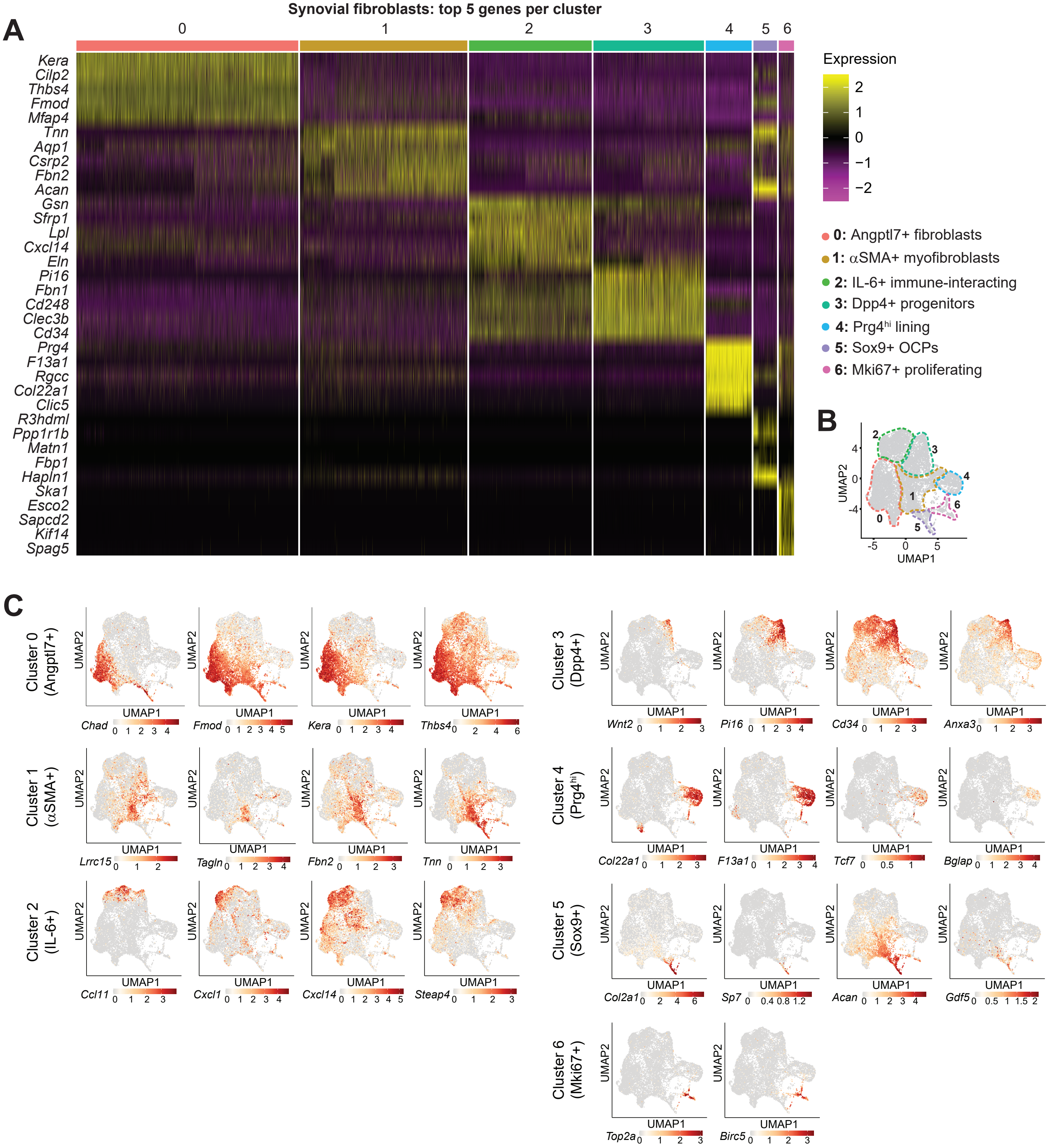

### Supplementary Figure 5

Fig S5. Functions and communication patterns of synovial fibroblast subsets

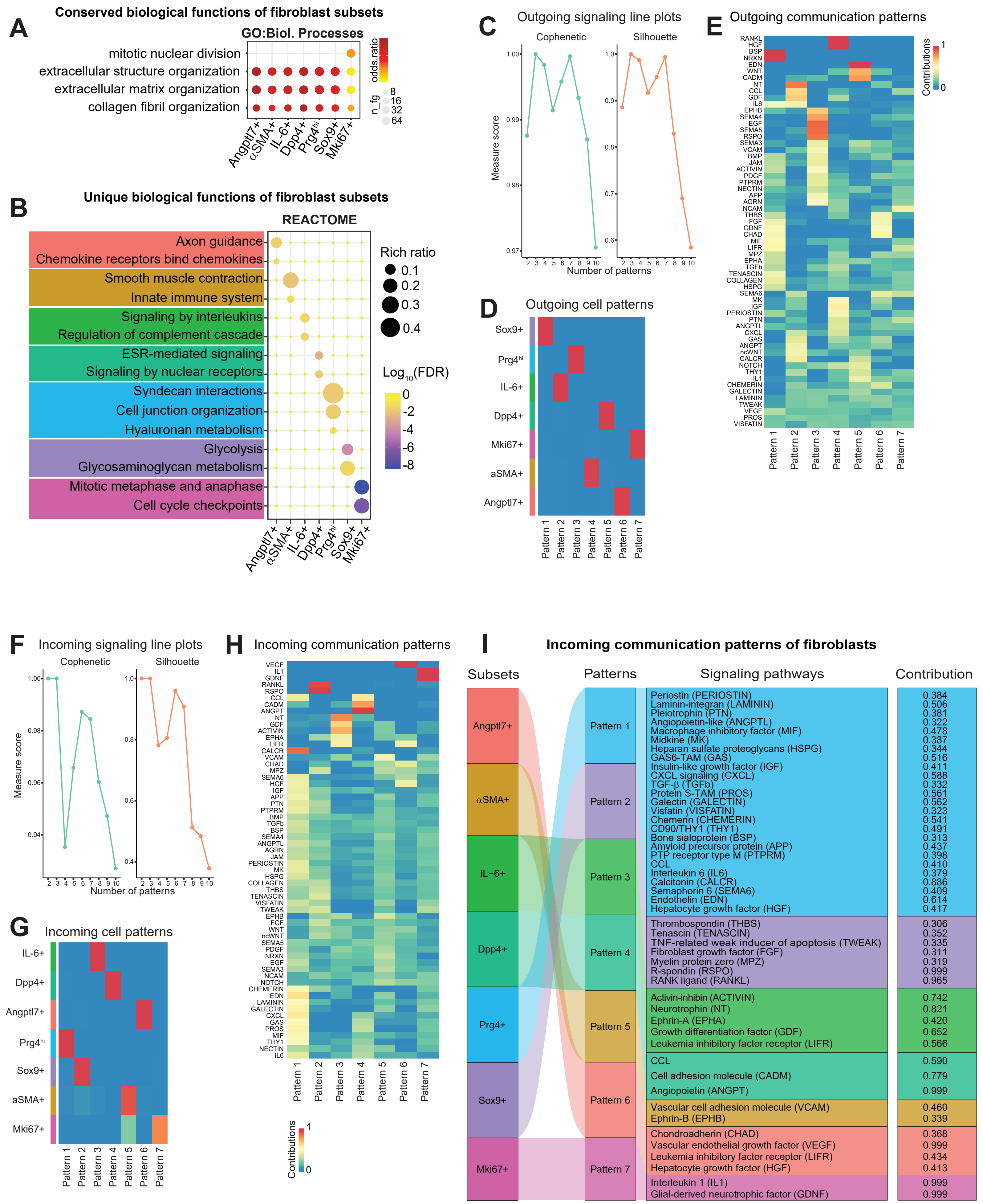

### Supplementary Figure 6

##### Fig S6. Wnt signaling is induced in synovium after joint injury

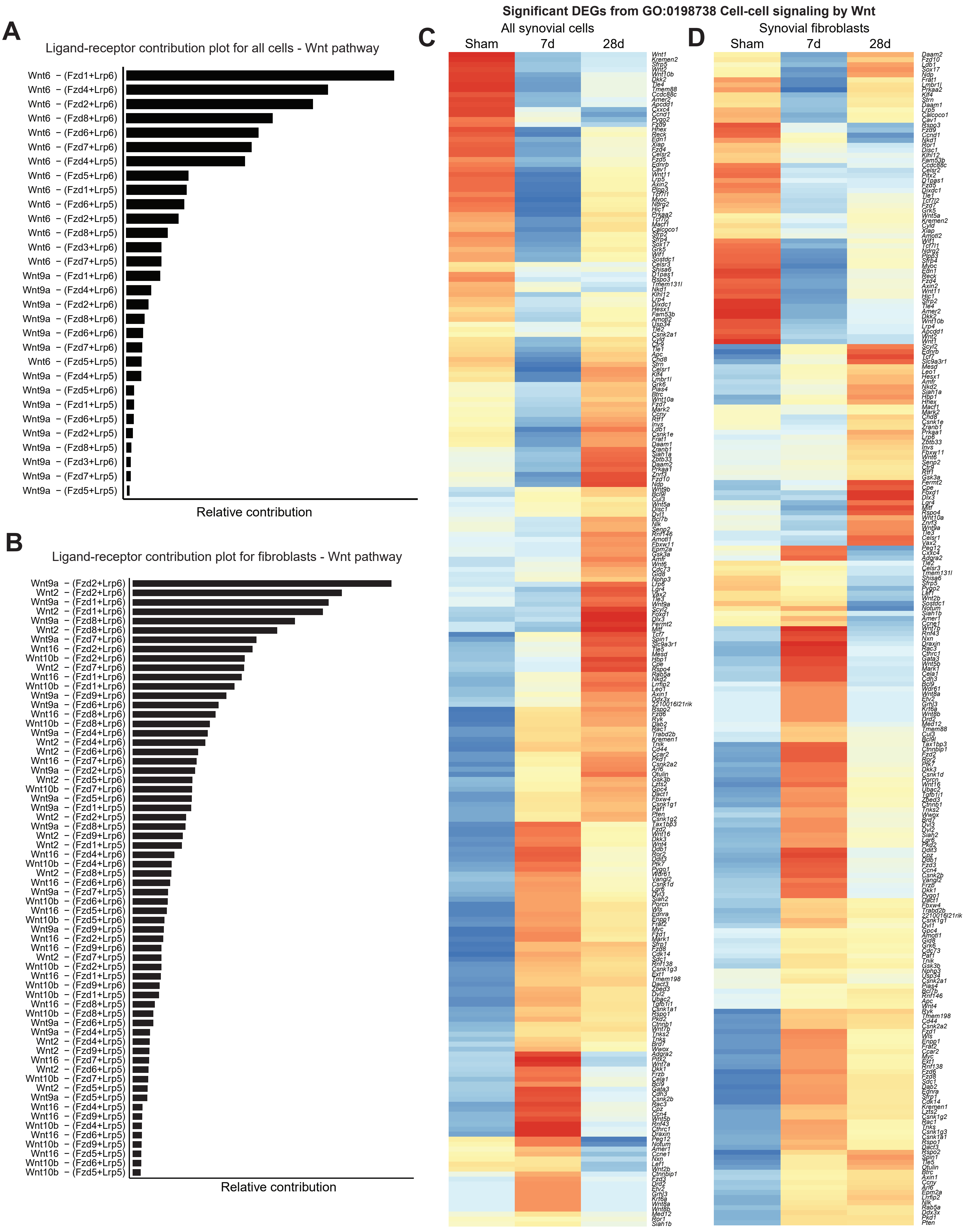

### Supplementary Figure 7

#### Fig S7. Flow cytometric analysis of synovium from injured Wnt-GFP reporter mice

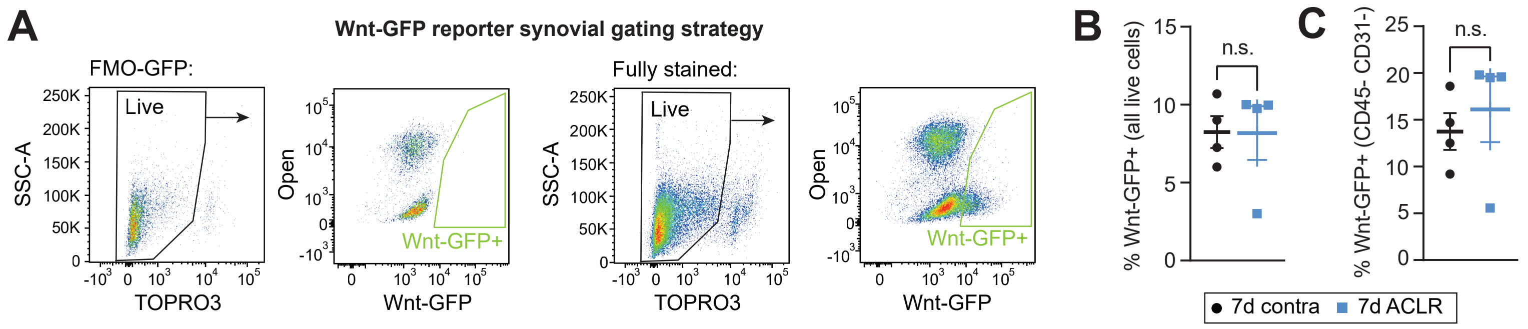

### Supplementary Figure 8

#### Fig S8. R-spondin 2 is activated in synovium following joint injury

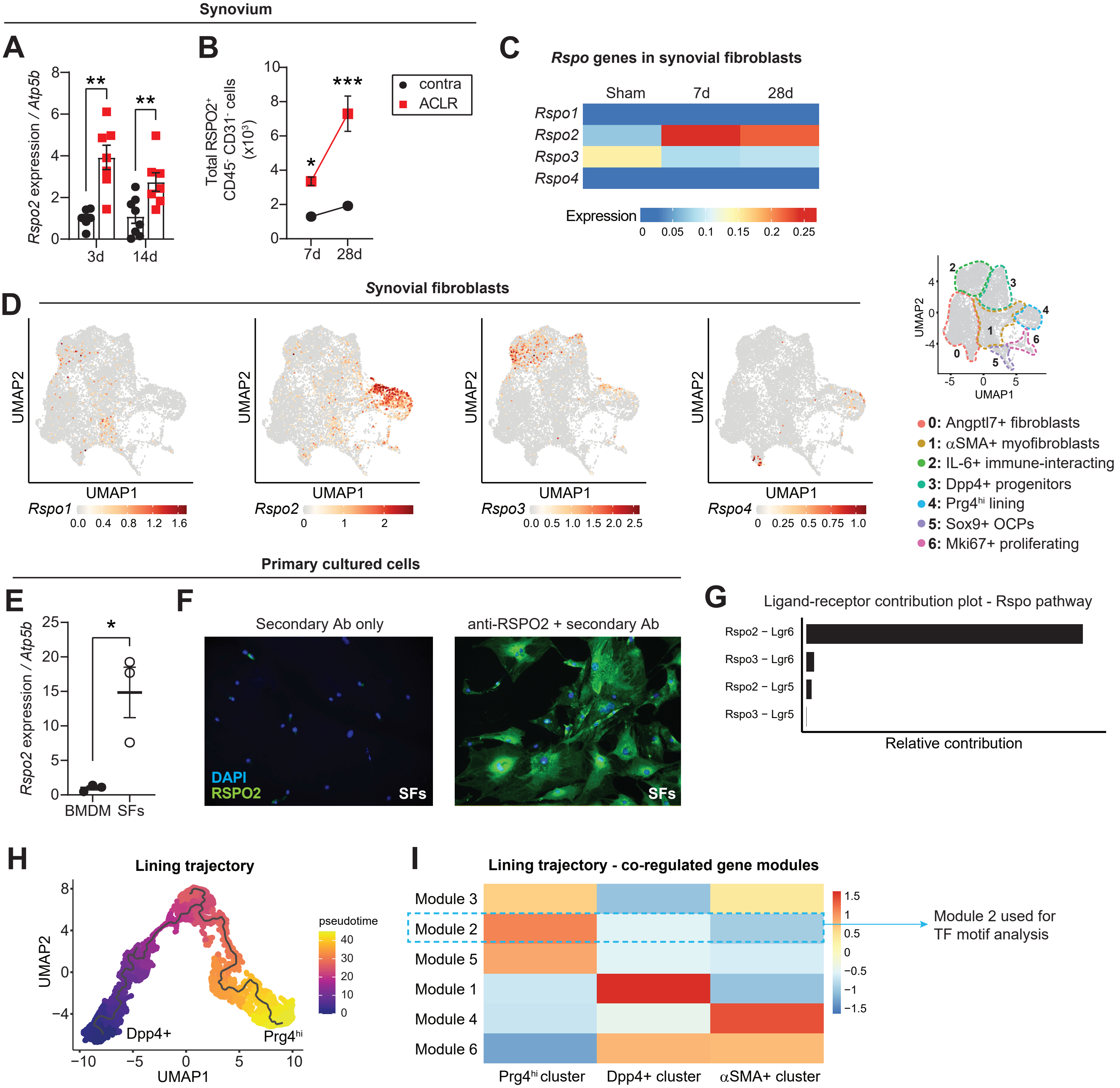

### Supplementary Figure 9

**Fig S9. R-spondin 2 orchestrates crosstalk between synovial fibroblasts and macrophages**

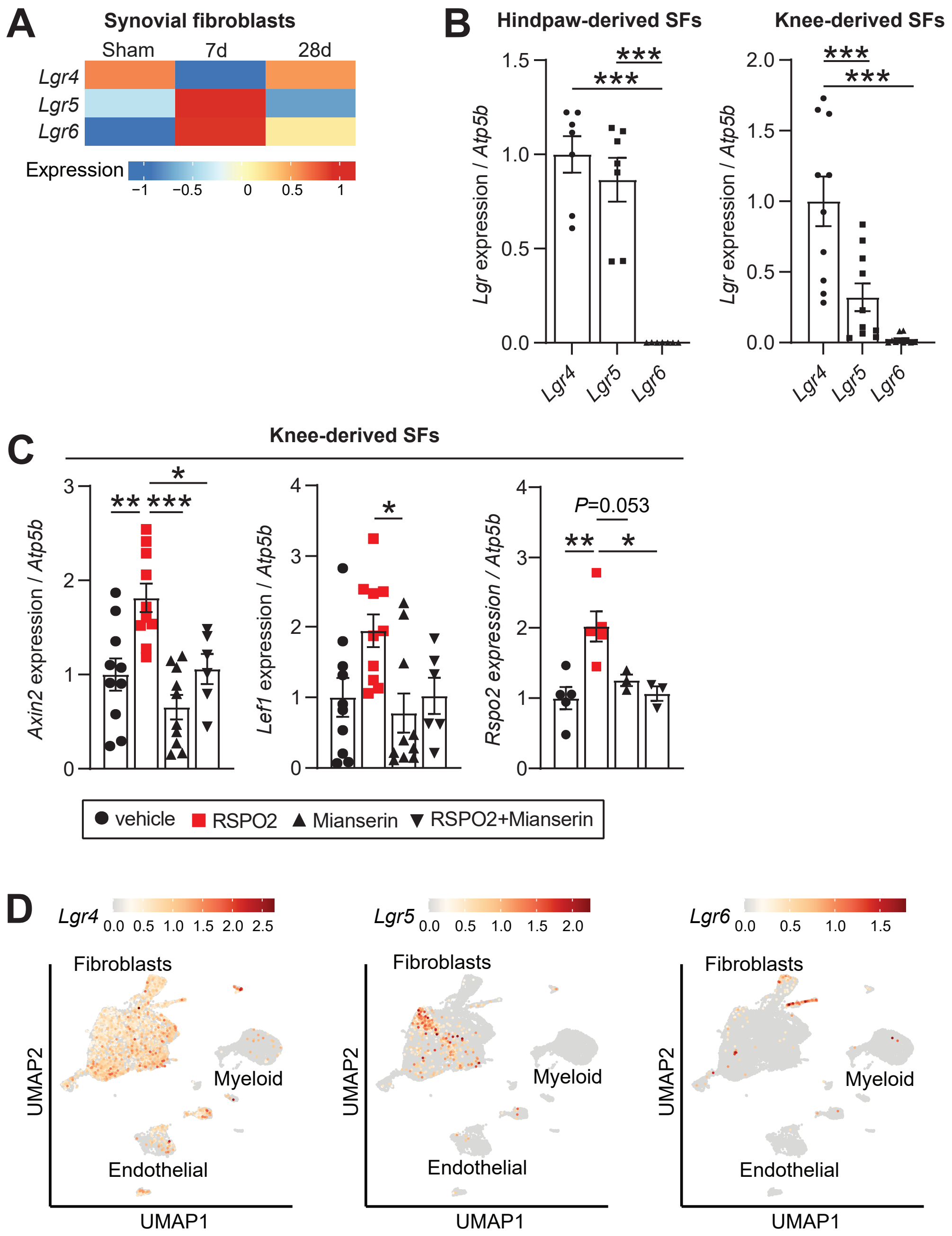

### Supplementary Figure 10

**Fig S10. R-spondin 2 orchestrates crosstalk between synovial fibroblasts and chondrocytes**

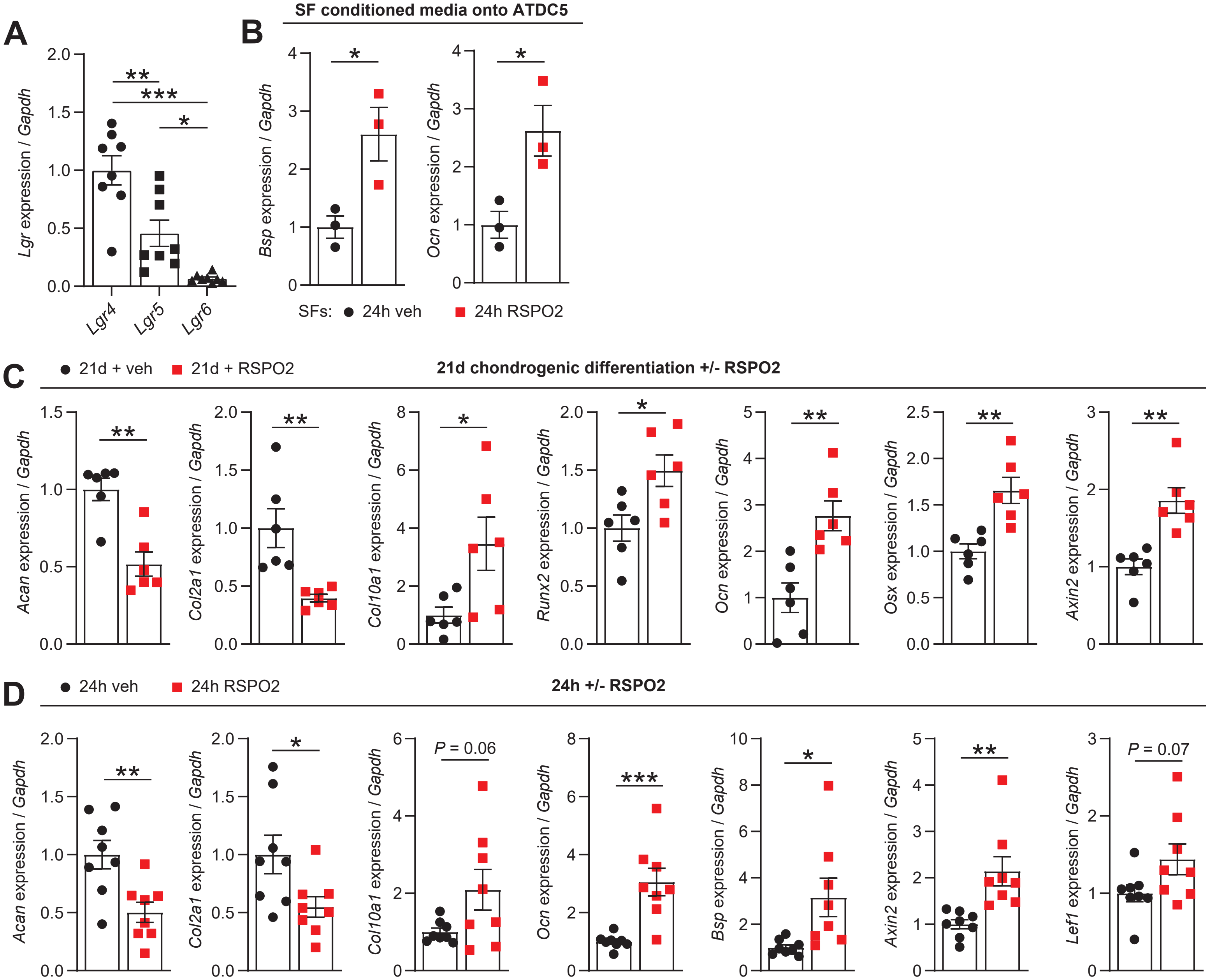
